# Multi-omics Profiling Identifies Molecular and Cellular Signatures of Regular Physical Activity in Human Peripheral Blood

**DOI:** 10.64898/2026.01.13.699221

**Authors:** Xuemei Song, Jingzhi Lv, Siliang Ge, Sihan Xu, Yisha Wu, Yuhui Zheng, Wenwen Zhou, Lifeng Li, Yan Zhang, Jiaqiang Zhang, Zhenxia Chen, Peng Gao, Pengbin Yin, Jianhua Yin, Chuanyu Liu

## Abstract

Regular physical activity is well established to protect against metabolic disorders and bolster immunity; yet, the underlying molecular and cellular mechanisms remain incompletely understood. We integrated plasma metabolomics and lipidomics with single-cell transcriptomic and chromatin accessibility profiles to decode the systemic impact of physical activity on human immunity and metabolism. Our data reveal that regular physical activity is linked to a coordinated metabolic signature marked by enhanced fatty acid oxidation and antioxidant defense. In circulating immune cells, regularly active individuals exhibited synchronous enhancement at both the chromatin accessibility and transcriptional levels for antigen presentation-related genes in antigen-presenting cells (APCs), particularly in classical monocytes, naive B cells, and switched memory B cells. Meanwhile, cytotoxic programs in CD8^+^ cytotoxic T and mature NK cells showed epigenetic pre-activation of effector function regulators. Intercellular communication analysis further revealed that regular exercise enhanced MHC-I/II signaling between APCs and T cells and suppressed inflammatory signaling networks. Together, these findings elucidate molecular mechanisms underlying the health benefits of regular exercise and offer a theoretical basis for enhancing public health and preventing chronic diseases.

## Introduction

Regular physical activity reduces the risk of chronic diseases(*1*), lowers cancer risk(*2*, *3*), while improving patient outcomes(*4*), enhances resistance to infection(*5*), delays aging(*6*), and mediated by circulating factors(*7*, *8*), ultimately integrating into whole-body benefits. However, existing mechanistic evidence is derived predominantly from human intervention studies or model organisms(*9–12*). Such controlled interventions differ substantially from free-living, spontaneously established regular physical activity habits. This discrepancy may lead to an incomplete understanding of the molecular and cellular mechanisms through which regular exercise confers clinical benefits, thereby limiting the external validity of current research findings.

Notably, metabolism and immunity are widely regarded as two central pillars through which exercise exerts its health-promoting effects(*13*, *14*). Metabolite and lipid profiling studies have identified candidate “exercise mimetics” or exercise-responsive metabolites, such as betaine, which has recently been characterized as an endogenous molecule triggered by sustained exercise that exerts systemic anti-inflammatory and neuroprotective effects(*10*); in parallel, flow cytometry and immunophenotyping studies have systematically revealed the remodeling effects of regular physical activity on the human immune system(*15*). Regular exercisers exhibit higher diversity of the naive and memory T-cell repertoire, a lower proportion of senescent T cells(*16*, *17*), a more anti-inflammatory phenotype of monocyte subsets(*18*, *19*), and enhanced functional adaptability, reduced senescence markers, and improved metabolic fitness in natural killer (NK) cells(*20*). Building on these advances, there is a growing need to move beyond isolated measurements toward a systems-level understanding of how regular physical activity remodels metabolic and immune networks. In particular, we still lack an integrated view of the heterogeneous responses within and across specific immune cell subsets, their dynamic interactions with one another, and how these processes are orchestrated through epigenetic regulation and metabolic reprogramming in response to exercise. Addressing these questions with integrated multi-omics approaches will be essential to construct a more holistic atlas of how regular physical activity reshapes immunometabolic circuits and, ultimately, to translate these insights into more precise and scalable health interventions.

Here, we conducted a comprehensive multi-omics study leveraging data from the Chinese Immune Multi-omics Cohort (CIMA)(*21*, *22*), a large-scale population-based cohort. By integrating plasma targeted metabolomics and lipidomics with single-cell transcriptomic and epigenomic analysis of peripheral blood mononuclear cells (PBMCs), we systematically mapped the molecular and cellular correlates of regular physical activity by comparing individuals with regular physical activity with sedentary controls. Our data reveal that regular physical activity is linked to a coordinated metabolic signature marked by enhanced fatty-acid oxidation and antioxidant defenses, together with widespread remodeling of immune cell states mediated by epigenetic reprogramming. Collectively, these results provide, to our knowledge, multi-dimensional atlas of molecular adaptations to regularly spontaneous exercise, spanning metabolism, chromatin regulation, and immunity. Beyond deepening mechanistic insight into how regular physical activity promotes health, this work highlights candidate biomarkers and regulatory pathways that may inform prevention and management of chronic diseases.

## Results

### Study Cohort and Physiological Characterization

Leveraging the CIMA natural cohort, we categorized participants based on self-reported physical activity (Figure 1A). Our classification criteria were informed by the WHO Guidelines on Physical Activity and Sedentary Behaviour(*23*). The regular exercise group (EX, n = 40) was defined as individuals who engaged in physical activity ≥ 3 times per week, accumulating ≥ 150 minutes of moderate-intensity or ≥ 75 minutes of vigorous-intensity activity. The sedentary control group (SED, n = 46) was defined as individuals who reported a weekly exercise duration of 0 minutes and identified “sitting” as their primary daily posture. The study cohort was characterized by a mean age of 28.2 years (SD ± 3.1) and body mass index of 21.3 kg/m² (SD ± 3.0) (Figure S1A). The two groups were well-balanced at baseline, with no statistically significant disparities observed in age, diet, smoking or alcohol status,or sleep duration(Figure S1A and Table S1). The SED group was predominantly female, consistent with global public health data showing higher physical inactivity rates among women(*24*, *25*). This sex imbalance in our cohort led to a significantly higher unadjusted mean BMI in EX. Given that sex and Body Mass Index (BMI) are key covariates for metabolic and immune phenotypes, subsequent analyses employed multiple linear regression models adjusted for sex and BMI to independently assess the effects of exercise.

**Figure 1.**
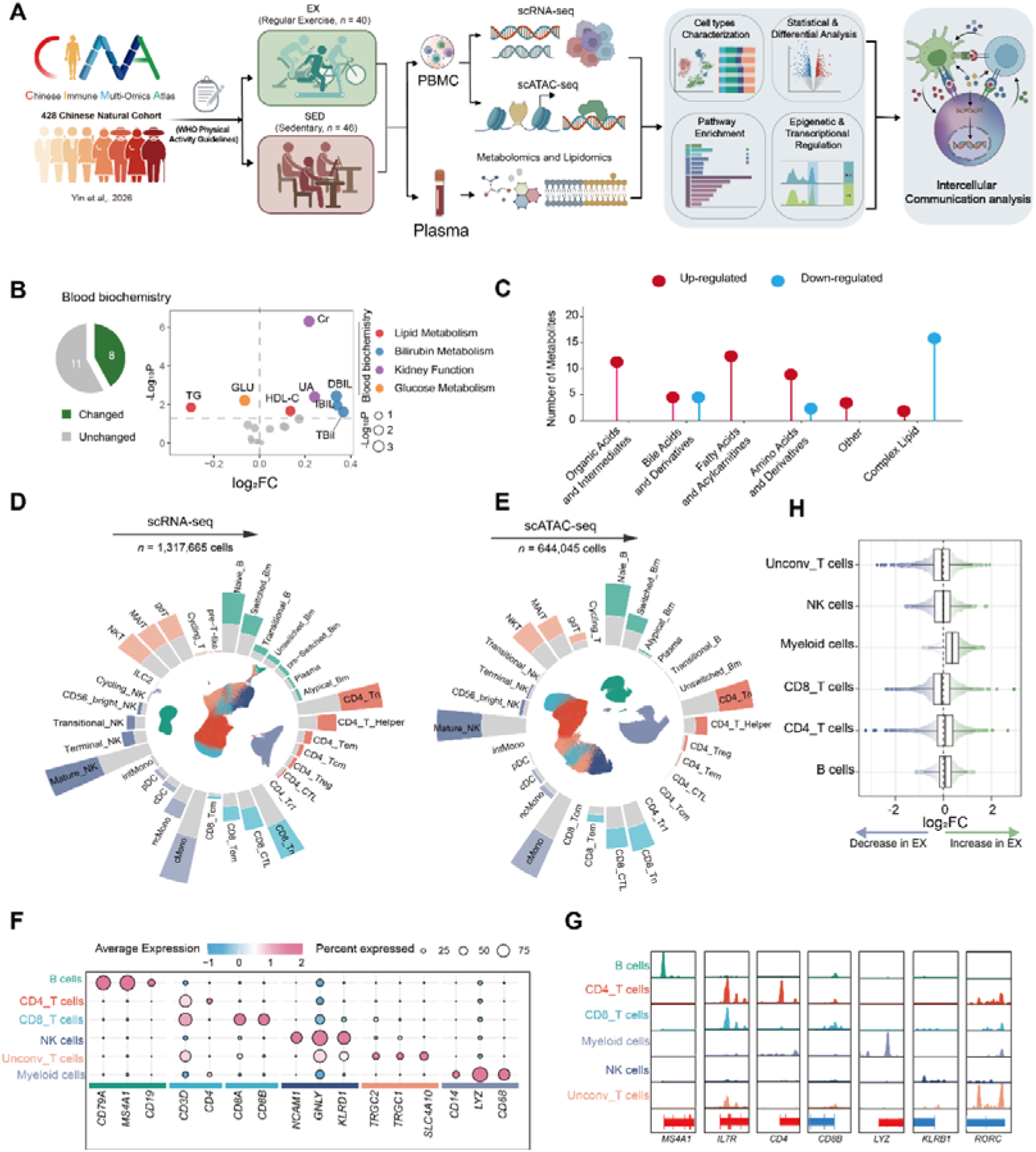
Study overview and multi-omics characterization of regular exercisers and sedentary controls. (A) Overview of the study design. A total of 86 participants from the CIMA cohort were classified into regular exercise (EX, n = 40) and sedentary (SED, n = 46) groups; all data were obtained from this cohort. (B) Left: pie chart showing the number of significantly changed (n = 8) and unchanged (n = 11) blood biochemical indices. Right: dot plot showing log_2_ fold change (log_2_FC) of blood biochemical indices in EX compared with SED, coloured by metabolic category. (C) Lollipop plot showing the number of significantly up-regulated (red) and down-regulated (blue) plasma metabolites and lipids in EX compared with SED, grouped by molecular class. (D) UMAP embedding and circular bar plot showing immune cell subtypes and their proportions identified by scRNA-seq (n = 1,317,665 cells), coloured by major lineage. (E) UMAP embedding and circular bar plot showing immune cell subtypes and their proportions identified by scATAC-seq (n = 644,045 cells), coloured by major lineage. (F) Dot plot showing the average expression (colour scale) and percentage of expressing cells (dot size) of canonical marker genes across major immune lineages. (G) Genome browser tracks showing chromatin accessibility at marker gene loci (*MS4A1*, *IL7R*, *CD4*, *CD8B*, *LYZ*, *KLRB1*, and *RORC*) across major immune cell types. (H) Beeswarmand and box plots showing the distribution of log_2_FC differences in cell neighborhoods across major immune cell lineages (EX vs. SED), as determined by Milo differential abundance analysis. Each point represents a cell neighborhood; blue and green indicate decrease and increase in EX, respectively. Box plots show median and interquartile range (IQR); whiskers extend to 1.5×IQR from the hinges.

To validate the physiological relevance of the classification based on exercise habits, we first compared routine blood biochemical markers. Of the 19 indices tested, eight differed significantly between EX and SED (Figure 1B). These alterations were reflected in glucose and lipid metabolism indicators: compared to SED, EX showed significantly lower fasting blood glucose (GLU) and triglyceride (TG) levels, along with significantly higher high-density lipoprotein cholesterol (HDL-C) levels (Figure S1B). This finding is consistent with a substantial body of prior research concluding that exercise confers significant cardiometabolic benefits(*26–28*). Beyond these classic signatures, EX showed significantly elevated levels of creatinine, uric acid, and bilirubin, all within normal clinical ranges (Figures 1B and S1C). Notably, creatinine is a product of muscle metabolism, and its elevation is directly associated with increased muscle mass and heightened metabolic turnover in exercise-trained individuals(*29*). Meanwhile, uric acid and bilirubin possess antioxidant properties, and their moderate elevation may reflect an enhancement in systemic antioxidant defense capacity(*30*, *31*).

In summary, classification based on exercise habits effectively captured distinct physiological states between EX and SED. These phenotypic data indicate that EX exhibits a multifaceted beneficial profile characterized by improved glucose and lipid metabolism, enhanced muscle metabolism, and augmented systemic antioxidant capacity.

### Multi-omics Profiling of Plasma and Circulating Immune Cells in Exercise versus Sedentary Individuals

To comprehensively elucidate the regulatory effects of regular exercise on systemic metabolism and immune features, we conducted a multi-omics analysis using data from 86 subjects in the CIMA cohort. This analysis integrated targeted plasma metabolomics, lipidomics, single-cell RNA sequencing (scRNA-seq), and single-cell assay for transposase-accessible chromatin sequencing (scATAC-seq) of PBMCs (Figure 1A). First, we performed metabolomic and lipidomic profiling on the plasma samples. Following standardized pre-processing, 321 metabolites and 718 lipid species were utilized for downstream analysis (Table S2). Comparative analysis revealed a distinct class-specific distribution pattern among the differential metabolites between EX and SED (Figure 1C). Specifically, metabolites associated with energy metabolism and substrate turnover (acylcarnitines and organic acids) were broadly upregulated in EX, whereas complex lipids were downregulated (Figure 1C and Table S2), indicating that regular exercise significantly remodels human metabolic program.

Subsequently, to investigate the impact of exercise on the immune system, we conducted scRNA-seq analysis on 1,317,665 high-quality cells from EX and SED (Figure 1D). Simultaneously, we performed an independent re-analysis and stringent filtering of the matched scATAC-seq raw fragments, ultimately retaining 644,045 high-quality cells (Figure 1E). Rigorous quality assessment ensured high data quality for both omics datasets (Figures S2A and S2B), with no significant batch effects detected between EX and SED (Figure S2C). Regarding cell type identification, we annotated the scRNA-seq data following the hierarchical annotation system of the CIMA cohort. The first level (L1) included six major immune cell types: myeloid cells, B cells, CD8^+^ T cells, CD4^+^ T cells, NK cells, and non-canonical T (Unconv_T) cells, which were further divided into 34 distinct subsets (Figures 1D, 1F and S2D). Subsequently, we annotated the scATAC-seq data using a cross-modal mapping strategy, achieving high concordance with the transcriptome at the L1 level and successfully resolving 30 subsets (Figure 1E). Gene activity scores based on chromatin accessibility aligned closely with transcriptomic cell type annotations (Figures S2E and S2F), with canonical marker gene promoters (*MS4A1*, *IL7R*, *CD4*, *CD8B*, *KLRB1* and *RORC*) exhibiting the expected lineage-specific accessibility patterns (Figure 1G). The proportions of major cell subsets remained stable between EX and SED in the scATAC-seq data (Figure S2G). In contrast, scRNA-seq revealed a significant increase in myeloid cells in EX (Figure S2G), a finding corroborated by Milo analysis(*32*) (Figure 1H). These findings indicate that regular exercise habits reshape the innate immune composition primarily at the transcriptomic level.

### Systemic Metabolic and Lipidomic Remodeling

To decipher the impact of regular physical activity on systemic metabolic homeostasis, we performed targeted metabolomics and lipidomics analyses on plasma samples from EX and SED. A total of 62 differentially abundant molecules were identified (Figure 2A). Specifically, 86% of the differential metabolites were upregulated in EX, including O-Adipoylcarnitine and p-Hydroxymandelic acid, while 94% of the differential lipids were downregulated, such as PG 18:1-18:2 and PC 18:1-20:4 (Figure 2A and Table S2). Pathway enrichment analysis revealed that the upregulated metabolites were significantly enriched in core energy metabolism pathways such as the tricarboxylic acid cycle (TCA cycle), as well as various amino acid conversion pathways involving phenylalanine/tyrosine metabolism and glycine, serine, and threonine metabolism, indicating more active energy metabolism and amino acid turnover in the exercising population (Figure 2B). Furthermore, pathways related to cofactor metabolism and glutathione metabolism, which is a key antioxidant defense system, were also significantly enriched.

**Figure 2.**
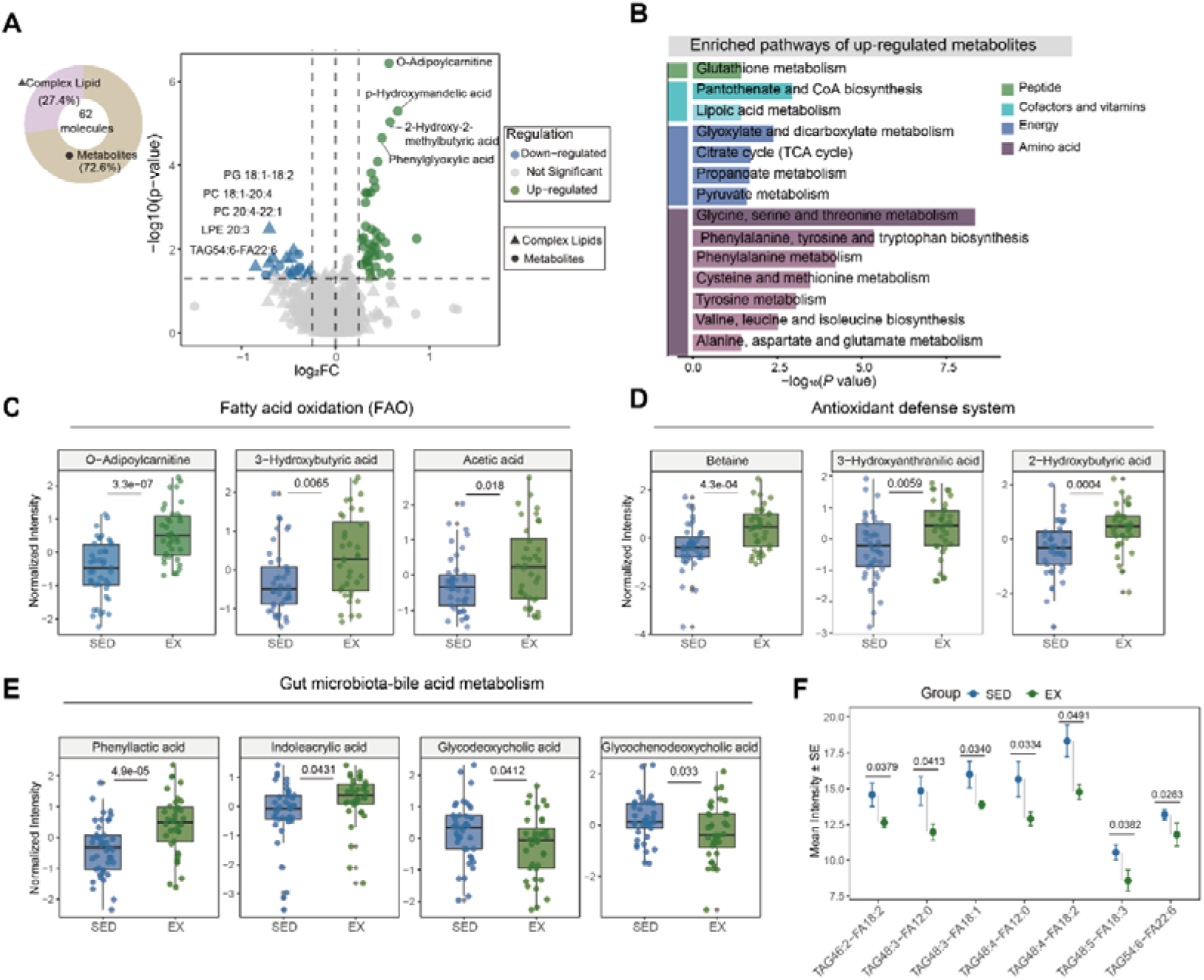
Plasma metabolomic and lipidomic remodeling in regular exercisers. (A) Volcano plot of differentially abundant plasma metabolites (circles) and complex lipids (triangles) in EX vs. SED. Pie chart inset shows the proportion of complex lipids and metabolites among the 62 significantly altered molecules. (B) Bar plot of enriched pathways of up-regulated metabolites in EX, coloured by pathway category. (C–E) Beeswarm and box plots showing normalized plasma levels of key metabolites related to fatty acid oxidation (C), antioxidant defense (D), and gut microbiota-bile acid metabolism (E) in SED and EX. (F) Dot plot showing mean intensity (± SE) of significantly altered triacylglycerol (TAG) species in SED and EX. *P-*values are indicated above each comparison. See also Table S2.

At the molecular level, we first focused on fatty acid oxidation (FAO), a central energy-yielding pathway during exercise (Figure 2C). The significant upregulation of O-adipoylcarnitine, a key acylcarnitine species reflecting mitochondrial fatty acid loading, indicates an increased flux of fatty acid transport into the mitochondria(*33*). Concurrently, elevated levels of 3-hydroxybutyrate, a primary ketone body, indicate the activation of hepatic fatty acid -oxidation and ketogenesis(*34*). Furthermore, the accumulation of acetic acid, a downstream product of fatty acid metabolism, further supports a comprehensive mobilization of FAO pathways across varying chain lengths(*35*). The coordinated elevation of these three core metabolites demonstrates that regular exercise systematically enhances the body’s fatty acid oxidative capacity.

Given that heightened energy metabolism is often accompanied by increased reactive oxygen species (ROS) production(*36*), we detailedly examined exercise-induced modulation of the antioxidant defense system (Figure 2D). In concert with elevated energy flux, several metabolites involved in redox homeostasis were synchronously upregulated in EX. Betaine, an important methyl donor, showed significantly higher plasma levels in EX. Notably, this molecule has recently been characterized as an endogenous exercise mimetic conferring anti-inflammatory and anti-aging protection(*10*). Furthermore, the concurrent elevation of 3-hydroxyanthranilic acid, which possesses radical-scavenging activity(*37*), and 2-hydroxybutyrate, an indirect marker of glutathione synthetic flux(*38*), collectively suggests that regular exercise reinforces a multi-layered endogenous antioxidant defense system. This trend was also observed for immunomodulatory metabolites, as exemplified by the increase in xanthurenic acid (another product of the tryptophan-kynurenine pathway) and γ-aminobutyric acid (GABA) (Table S2). These molecules can regulate the function and inflammatory status of immune cells such as T cells and monocytes/macrophages through the aryl hydrocarbon receptor or GABAergic signaling pathways(*39*, *40*).

Furthermore, we observed that regular exercise also reshapes gut microbiota-related metabolites (Figure 2E). Levels of microbially derived aromatic amino acid metabolites—phenyllactic acid and indoleacrylic acid—were significantly elevated in EX. Indoleacrylic acid has been shown to enhance intestinal barrier function and modulate innate immunity, thereby contributing to reduced systemic inflammation(*41*). In parallel, levels of two secondary bile acids, glycodeoxycholic acid and glycochenodeoxycholic acid, were reduced in EX. Bile acids are signaling molecules that regulate intestinal and hepatic immunity(*42*), and the altered levels of these secondary bile acids may influence local and systemic immune activity.

At the lipid level, a broad downregulation was observed in EX. Triacylglycerols (TAG), the primary energy-storage form, exhibited widespread reduction (Figure 2F), consistent with the well-established effect of exercise-promoted lipolysis and fatty acid utilization(*43*). Multiple membrane phospholipid subclasses, including lysophosphatidylethanolamine (LPE), phosphatidylcholine (PC), phosphatidylglycerol (PG), and phosphatidylinositol (PI), also showed decreased abundance (Figure S3A), reflecting extensive remodeling of the lipid metabolic network. Notably, distinct species within the same phosphatidylethanolamine (PE) subclass displayed opposite abundance changes: while PE 20:4−22:5 was downregulated, PE 20:4−22:1 was upregulated in EX (Figure S3B). Overall, these lipid alterations, together with enhanced fatty acid oxidative capacity, constitute a characteristic metabolic adaptation to regular exercise.

### Myeloid Cells Exhibit Enhanced Antigen Presentation Following Regular Exercise

Given the pivotal role of myeloid cells in sensing metabolic signals and regulating inflammation and antigen presentation(*44*, *45*), elucidating how regular physical activity remodels these cell populations is essential. Myeloid cells were stratified into five subsets using canonical markers: classical monocytes (cMono), intermediate monocytes (intMono), non-classical monocytes (ncMono), classical dendritic cells (cDC), and plasmacytoid dendritic cells (pDC) (Figures 3A, S4A, and S4F). Conventional cell proportion analysis showed no significant differences (Figure S5A). Differential abundance testing with the Milo analysis at the local neighborhood level revealed significant expansions of intMono, cMono, and ncMono in EX (median log_2_FC: +0.37, +0.31, +0.25; Figure 3B). This suggests that regular physical activity remodels circulating myeloid cells, with a pronounced effect on the composition of monocyte subsets.

**Figure 3.**
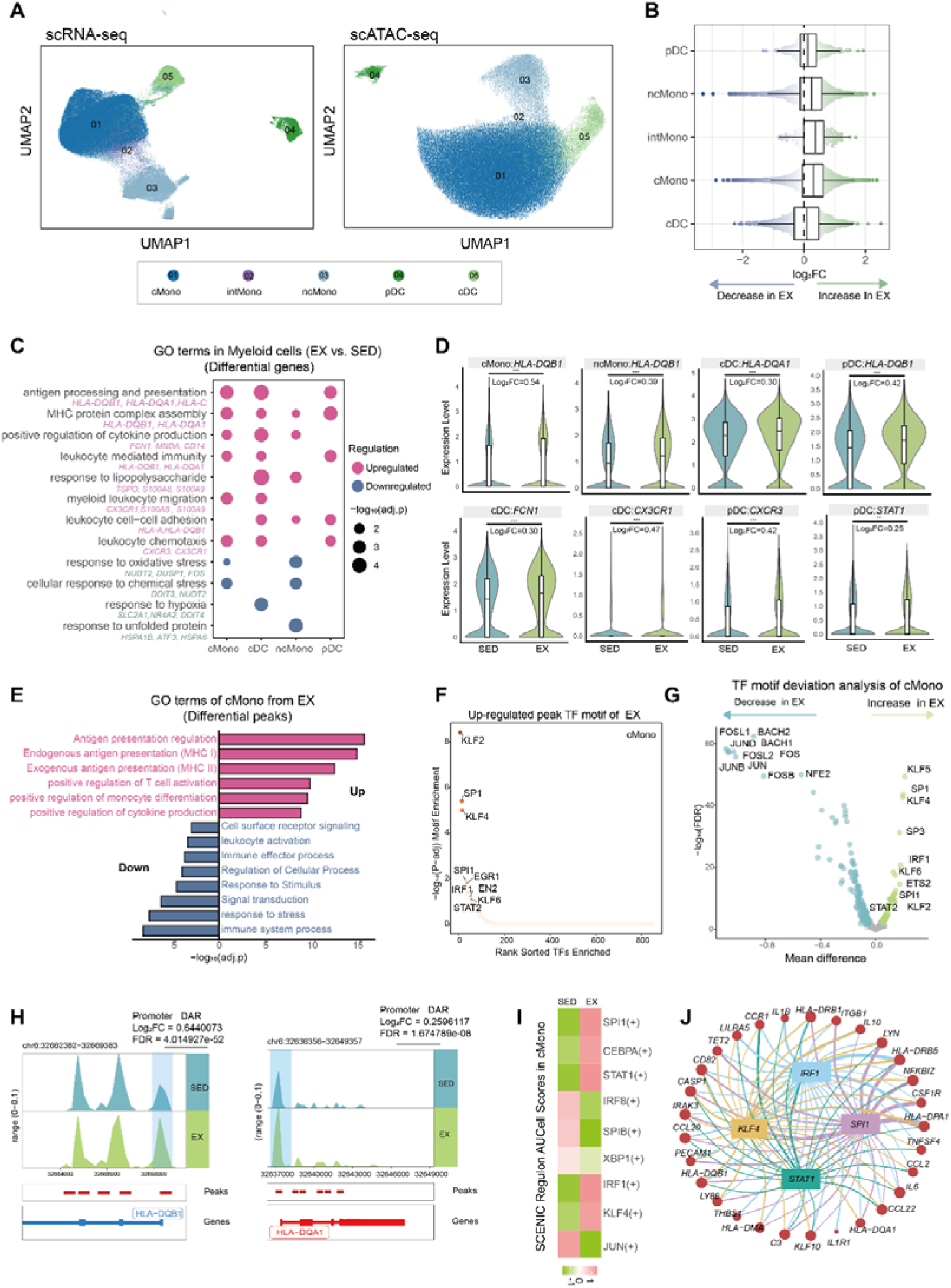
Epigenetic and transcriptional characteristics of myeloid cells in regularly active individuals. (A) UMAP visualization of myeloid cell subsets identified by scRNA-seq (left) and scATAC-seq (right), colored by cell type. (B) Beeswarmand and box plots showing the distribution of log_2_FC differences in cell neighborhoods across myeloid cell subsets (EX vs. SED). Box plots are created in a similar fashion as in Figure 1H. (C) Bubble plot of Gene Ontology (GO) enrichment analysis for differentially expressed genes (DEGs) in myeloid subsets (EX vs. SED). Dot size represents -log_10_(adjusted *p*-value); color indicates regulation direction (pink: upregulated in EX; blue: downregulated in EX). (D) Violin plots overlaid with box plots showing expression levels of selected genes associated with antigen presentation, innate immunity, and cellular migration across myeloid subsets in SED and EX. Log_2_FC and adjusted p-values are indicated. \*\**p* < 0.01, \*\*\**p* < 0.001. (E) GO enrichment analysis of differentially accessible regions (DARs) in cMono (EX vs. SED). Bars extending right (pink) indicate pathways enriched in open chromatin regions in EX; bars extending left (blue) indicate decreased accessibility in EX. Bar length represents –log_10_(*p*-value). (F) Transcription factor (TF) motif enrichment analysis of differentially accessible peaks with increased accessibility in cMono of EX. TFs are ranked by statistical significance; top-ranked motifs are labeled. (G) TF motif deviation analysis in cMono. Each point represents a TF motif; x-axis shows mean chromatin accessibility difference (EX vs. SED); y-axis shows –log_10_(FDR). Green and blue points indicate TF motifs with significantly increased and decreased accessibility in EX, respectively; gray points indicate non-significant motifs. (H) Genome browser tracks of chromatin accessibility at the *HLA-DQB1* (left) and *HLA-DQA1* (right) loci in cMono from SED and EX. Promoter DARs are highlighted; log_2_FC and FDR values are indicated. (I) Heatmap showing the scaled mean regulon areas under the curve per cell (AUCell) scores of selected transcription factors in cMono between SED and EX. Red and green indicate higher and lower regulon activity, respectively. (J) Network diagram of the core transcriptional regulatory module in cMono, centered on *SPI1*, *IRF1*, *KLF4*, and *STAT1*. Nodes represent target genes involved in antigen presentation and inflammatory/chemotactic regulation; edge colors indicate regulatory relationships.

To dissect the functional alterations of myeloid subsets induced by regular exercise, we performed differentially expressed gene (DEG) and pathway enrichment analyses (Table S3). Gene Ontology (GO) analysis of upregulated genes in EX revealed enrichment in antigen processing and presentation, major histocompatibility complex (MHC) protein complex assembly, and leukocyte chemotaxis, whereas downregulated DEGs were predominantly linked to oxidative stress and hypoxia responses (Figure 3C). Pathway scoring analysis further corroborated these shifts, manifesting as enhanced antigen presentation capabilities alongside a systemic attenuation of inflammatory and stress-related signatures across multiple subsets in EX (Figure S5B). Consistent with their enhanced functional state, the core antigen presentation gene *HLA-DQB1* showed pronounced elevation in cMono, ncMono, and pDC, while *HLA-DQA1* and the innate immune sensor *FCN1* were markedly upregulated in cDC (Figure 3D). Furthermore, pDC exhibited increased expression of the key transcriptional regulator *STAT1*, alongside elevated levels of migration-associated receptors, including *CX3CR1* in cDC and *CXCR3* in pDC, suggesting a concurrent enhancement of antigen presentation and cellular migration capacity in myeloid cells.

To investigate the epigenetic mechanisms underpinning these functional changes, we compared the chromatin accessibility of myeloid cell subsets between EX and SED using scATAC-seq. Analysis of differentially accessible regions (DARs) revealed that 94.3% of DARs across all myeloid subsets were specifically enriched in cMono (Figure S5C and Table S4). GO enrichment analysis of DARs in cMono showed that regions with increased accessibility were enriched for pathways such as antigen presentation and T cell activation, while regions with decreased accessibility were associated with stress response pathways (Figure 3E). Transcription factor (TF) motif enrichment and motif deviation analysis characterized a distinct regulatory signature in EX (Figures 3F and 3G). Specifically, regions with increased accessibility in EX were primarily enriched for TFs supporting antigen presentation (*IRF1* and *SPI1*)(*46*, *47*), viral defense (*STAT2*)(*48*), and immune homeostasis and migration (*KLF2* and *KLF4*)(*49*, *50*). Conversely, the accessibility of motifs for regulatory factors mediating stress and inflammatory responses, such as the AP-1 family (*JUN, JUNB, FOS* and *FOSB*) and *BACH1*, was significantly reduced. To determine the epigenetic basis for enhanced antigen presentation, we examined chromatin accessibility at loci linked to this function in cMono. Analysis showed that the promoter regions of multiple key antigen presentation genes (*HLA-DQA1* and *HLA-DQB1*) exhibited significantly enhanced chromatin accessibility in EX (Figure 3H), consistent with their upregulated mRNA expression in this subset (Figure 3D). Additionally, EX exhibited reduced chromatin accessibility at the promoters of *CXCL8* and *CCL7* and a distal regulatory element associated with *CCL2* (Figure S5D). The reduced promoter accessibility of *CXCL8* was consistent with its downregulated mRNA expression (Figure S5E).

Next, we assessed regulon factor activity in cMono at the transcriptional level using the Single-Cell Regulatory Network Inference and Clustering (SCENIC) algorithm(*51*). Results demonstrated that the activities of transcription factors such as *SPI1*, *STAT1*, *IRF1*, *and KLF4* were significantly higher in EX than in SED (Figure 3I), highly consistent with the scATAC-seq data indicating upregulated accessibility of the corresponding motifs (Figures 3F and 3G). To understand how key transcription factors cooperatively regulate cMono function, we identified a core regulatory module driven by *SPI1*, *IRF1*, *KLF4*, *and STAT1* (Figure 3J). This core hub orchestrates diverse immune programs, spanning antigen presentation (*HLA-DQA1* and *HLA-DQB1*) and the regulation of inflammation and chemotaxis (*IL1B*, *IL10*, *CCL20*, and *CCL2*). In summary, regular physical activity remodels the epigenetic landscape and transcriptional profiles of myeloid cells. Collectively, EX exhibits enhanced antigen presentation and interferon production alongside altered chemotaxis, with the most pronounced synergistic effects observed in cMono.

### Epigenetic Activation of Effector Functions in T Cells by Regular Exercise

Previous studies have shown that regular exercise can modulate the composition and functional status of circulating T cells(*14*, *52*), how exercise drives these functional changes via epigenetic reprogramming at the single-cell level remains to be systematically elucidated. Using classical marker genes, we identified 17 T cell subsets, consisting of ten CD8^+^ and unconv_T cell subsets: CD8^+^ naive T cells (CD8^+^ Tn), CD8^+^ central memory T cells (CD8^+^ Tcm), CD8^+^ effector memory T cells (CD8^+^ Tem), CD8^+^ cytotoxic T cells (CD8^+^ CTL), mucosal-associated invariant T cells (MAIT), γδT1, γδT2, proliferating T, and pre-T-like cells (Figures 4A, S4B and S4G). Additionally, we identified seven CD4^+^ T cell subsets (Figures S7A, S4C and S4H). Proportional analysis revealed that overall compositions of CD8^+^ and unconv_T cells remained stable between EX and SED (Figure S6A). However, Milo analysis identified a significant expansion of pre-T-like cells in EX (Figure 4B). Within the CD4^+^ T cell compartment, EX exhibited increased Treg proportions at the epigenetic level (Figure S7B), accompanied by a reduction in CD4^+^ cytotoxic T lymphocytes (CD4^+^ CTLs) (Figure S7C).

**Figure 4.**
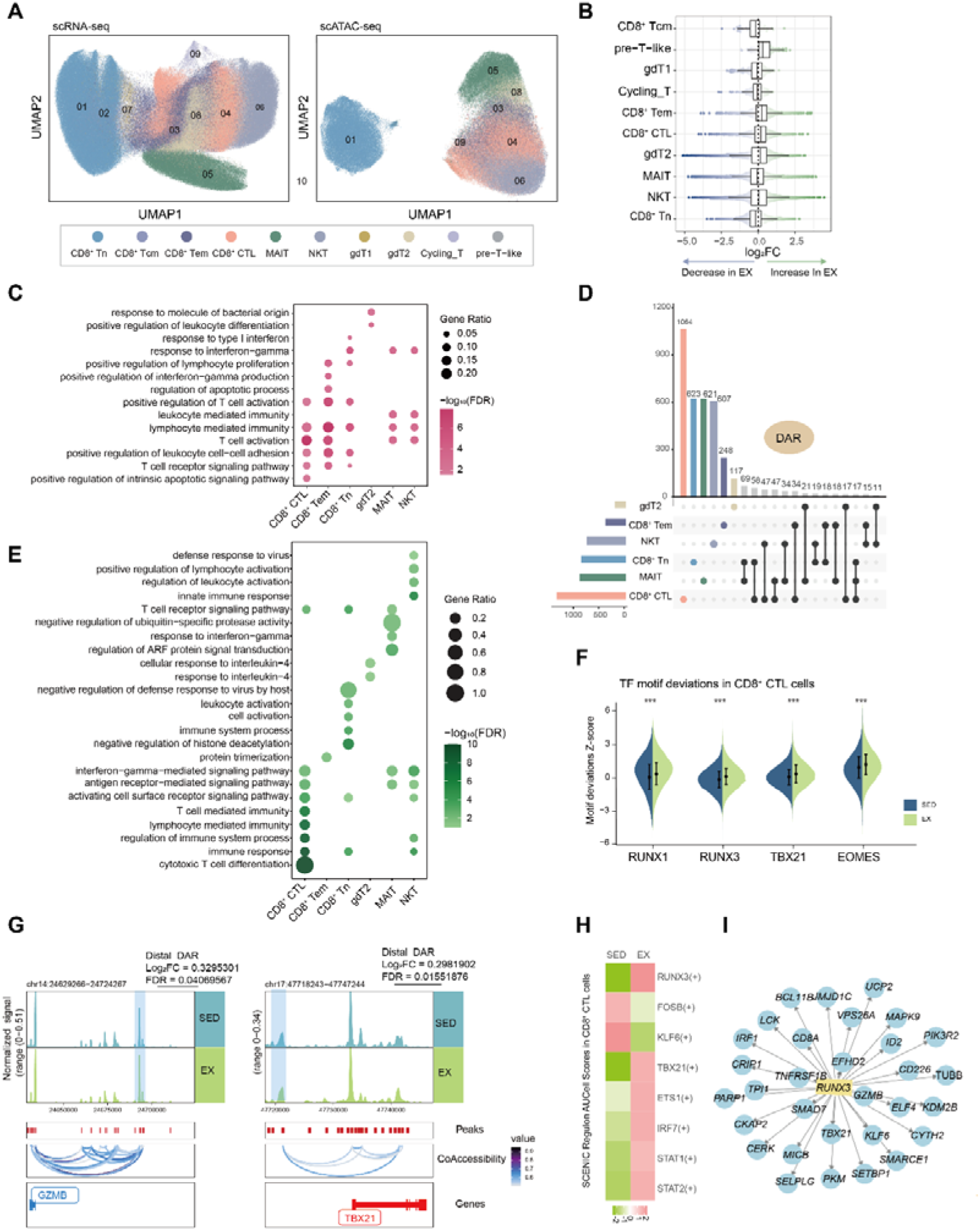
Epigenetic and transcriptional signatures of T cells associated with regular exercise. (A) UMAP visualization of CD8^+^ and unconventional T cell subsets identified by scRNA-seq (left) and scATAC-seq (right), colored by cell type. (B) Beeswarmand and box plots showing the distribution of log_2_FC differences in cell neighborhoods across CD8^+^ and unconventional T cell subsets (EX vs. SED). Box plots are created in a similar fashion as in Figure 1H. (C) Bubble plot of GO enrichment analysis for upregulated DEGs in CD8L and unconventional T cell subsets (EX vs. SED). Dot size represents gene ratio; color indicates –log_10_(FDR). (D) UpSet plot showing the number and overlap of upregulated DARs across CD8^+^ and unconventional T cell subsets in EX. Bar height indicates the number of DARs per subset or intersection. (E) Bubble plot of GO enrichment analysis for upregulated DARs across CD8^+^ and unconventional T cell subsets in EX. Dot size represents gene ratio; color indicates –log_10_(FDR). (F) Violin plots showing TF motif deviation Z-scores for *RUNX1*, *RUNX3*, *TBX21*, and *EOMES* in CD8^+^ CTL cells from SED and EX. Adjusted *p*-values are indicated. \*\*\**p* < 0.001. (G) Genome browser tracks of chromatin accessibility at distal regulatory regions of *GZMB* (left) and *TBX21* (right) loci in CD8^+^ CTL cells from SED and EX. Co-accessibility links and distal DARs are highlighted; log_2_FC and FDR values are indicated. (H) Heatmap showing the scaled mean regulon AUCell scores of selected transcription factors in CD8^+^ CTL cells between SED and EX. Red and green indicate higher and lower regulon activity, respectively. Network diagram of the *RUNX3* downstream transcriptional regulatory module in CD8^+^ CTL cells.

We identified DEGs and performed functional enrichment analysis (Table S3), broad transcriptomic alterations were present across these CD8^+^ and unconv_T cells (Figure S6B). GO enrichment analysis of upregulated genes in EX revealed significant enrichment of pathways related to T cell activation, T cell receptor (TCR) signaling, IFN-γ response, and lymphocyte-mediated immunity (Figure 4C). Within the CD4^+^ compartment, transcriptional changes were most pronounced in CD4^+^ Tn and T Helper cells (Figure S7D), with upregulated genes enriched in T cell activation, IL-12 production, and type I interferon signaling (Figures S7E and S7F). Pathway activity scoring further corroborated these functional shifts, revealing significantly elevated scores in EX across multiple T cell subsets, including CD8^+^ CTL, CD8^+^ Tem, CD8^+^ Tn, NKT, MAIT, and γδT1 cells, for processes related to T cell-mediated immunity, cytotoxicity, T cell activation, and type II interferon production (Figure S6C). Notably, the expression of key cytotoxic genes (*GZMH*, *GZMA* and *PRF1*) was significantly increased in CD8^+^ CTL, CD8^+^ Tem, and NKT cells (Figure S6D). Overall, the transcriptomic features collectively indicate a systemic shift in T cell function, characterized by a broad upregulation of activation programs and a reinforcement of cytotoxic effector.

To determine how regular exercise drives functional adaptation in T cells through epigenetic reprogramming, we performed DAR analysis across all subsets (Table S4). CD8^+^ CTLs exhibited the highest number of upregulated DARs (Figure 4D), with multiple DARs shared between two or more subsets. In CD8^+^ CTLs, upregulated DARs were enriched for pathways related to cytotoxic T cell differentiation, TCR signaling, and lymphocyte-mediated immunity. In CD8^+^ Tn cells, enrichment was observed for pathways involved in T cell activation and immune response regulation. Pathways associated with IFN-γ signaling, antigen receptor-mediated signaling, and TCR signaling were commonly enriched in CD8^+^ CTLs, MAIT cells, and NKT cells. Additionally, NKT cells showed enrichment for innate immune response and positive regulation of lymphocyte activation (Figure 4E).

To resolve the transcriptional regulatory basis of these functional changes, we performed motif deviation analysis based on scATAC-seq data. In CD8^+^ CTLs, the upregulated DARs were significantly enriched for binding motifs of effector function master regulators, including *EOMES*, *RUNX3*, *RUNX1*, and *TBX21* (Figures 4F and S6E). *TBX21* (T-bet) and *EOMES* are master regulators driving CD8^+^ T cell differentiation into effector cells, and their upregulation is directly linked to enhanced cytotoxic function(*53*, *54*). Similarly, motif deviation analysis identified increased accessibility for key transcription factor binding motifs in other subsets. These included motifs for *RORC*, *RUNX3*, *STAT1*, and *STAT4* in MAIT cells (Figure S6F), and for *LEF1*, *TCF7*, and *RUNX3* in CD8^+^ Tn cells (Figure S6G). Notably, chromatin accessibility was significantly increased at distal regulatory regions of the effector genes *TBX21* and *GZMB* in CD8^+^ CTLs (Figure 4G). Similarly, accessibility was elevated at the promoter of *IRF1* in MAIT cells (Figure S6H) and within an intron of *TCF7* in CD8^+^ Tn cells (Figure S6I). SCENIC analysis confirmed elevated regulatory activity of key transcription factors, including *RUNX3* and *TBX21*, in CD8^+^ CTLs of EX (Figure 4H). Given that *RUNX3* showed the most significant change across both omics layers, we constructed its downstream regulatory network (Figure 4I), which confirmed that *RUNX3* directly targets key effector genes, including *GZMB*, *TBX21*, and *IRF1*.

In summary, single-cell multi-omics data reveal a global remodeling of T cell function with regular exercise: it not only enhances the cytotoxic execution capacity of effector cells, with CD8^+^ CTLs at the core, but also broadly augments the signal responsiveness and activation preparedness of naive and non-canonical subsets.

### Functional Reprogramming of the NK Cell Epigenome and Transcriptome by Regular Exercise

As key innate effector cells, NK cells are pivotal in antiviral defense, tumor surveillance, and bridging innate and adaptive immunity(*55*). However, the molecular mechanisms by which regular exercise enhances NK cell function remain incompletely understood. We first classified NK cells into six subsets based on classical marker genes: mature NK cells, transitional NK cells, type 2 innate lymphoid cells (ILC2), CD56^bright^ NK cells, terminal NK cells, and cycling NK cells (Figures 5A, S4D, and S4I). Notably, the proportion of CD56^bright^ NK cells was significantly reduced in EX (Figure S8A), consistent with the results obtained from the Milo algorithm (Figure 5B).

**Figure 5.**
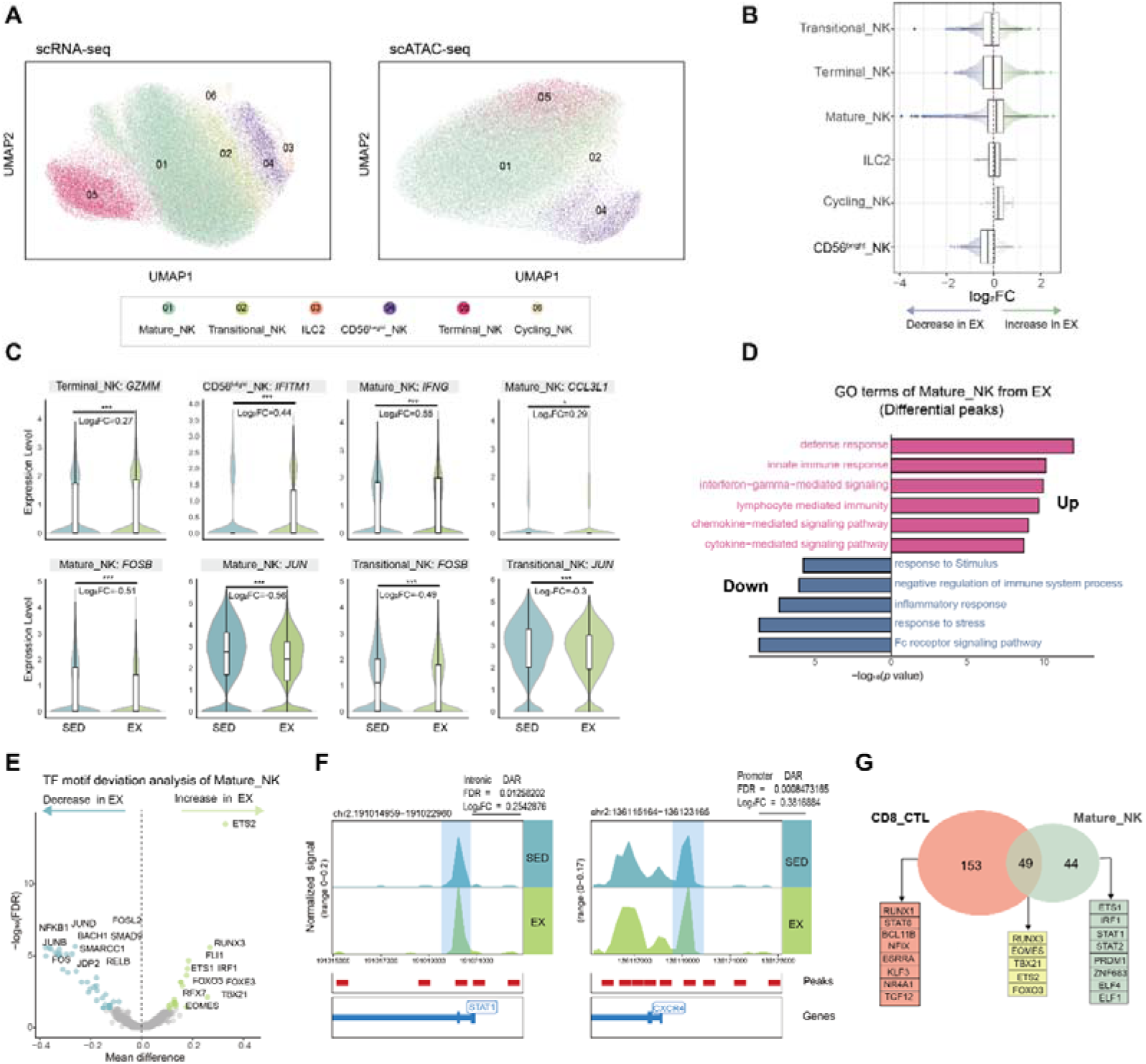
Epigenetic and transcriptional profiles of NK cells associated with regular exercise. (A) UMAP visualization of NK cell subsets identified by scRNA-seq (left) and scATAC-seq (right), colored by cell type. (B) Beeswarmand and box plots showing the distribution of log_2_FC differences in cell neighborhoods across NK cell subsets (EX vs. SED), as determined by Milo differential abundance analysis. Box plots are created in a similar fashion as in Figure 1H. (C) Violin plots overlaid with box plots showing expression levels of selected genes across NK cell subsets in SED and EX. Log_2_FC adjusted *p*-values and statistical significance are indicated. \**p* < 0.05, \*\*\**p* < 0.001. (D) GO enrichment analysis of DARs in Mature_NK cells (EX vs. SED). Bars extending right (pink) indicate pathways enriched in open chromatin regions in EX; bars extending left (blue) indicate decreased accessibility in EX. Bar length represents –log_10_(*p*-value). (E) TF motif deviation analysis in Mature_NK cells. Each point represents a TF motif; x-axis shows mean chromatin accessibility difference (EX vs. SED); y-axis shows –log_10_(FDR). Green and blue points indicate TF motifs with significantly increased and decreased accessibility in EX, respectively; gray points indicate non-significant motifs. (F) Genome browser tracks of chromatin accessibility at the intronic DAR of *STAT1* (left) and promoter DAR of *CXCR4* (right) in Mature_NK cells from SED and EX. Log_2_FC and FDR values are indicated. (G) Venn diagram showing the overlap of significantly upregulated TFs between CD8^+^ CTL and Mature_NK cells in EX. Shared and cell type-specific TFs are listed.

To evaluate functional changes in NK cells, we identified DEGs (Table S3) and performed pathway scoring analysis (Figure S8B). Elevated scores for NK cell-mediated cytotoxicity, NK cell activation, and IFN-γ response were observed across multiple NK subsets in EX, whereas scores for inflammatory response and cellular stress response were generally reduced (Figure S8B). In EX, subset-specific gene expression changes were observed in NK cells. *GZMM* was upregulated in terminal NK cells, *IFITM1* in CD56^bright^ NK cells, and *IFNG* and *CCL3L1* in mature NK cells. In contrast, *FOSB* and *JUN* were downregulated in both transitional and mature NK subsets (Figure 5C).

Epigenetically, we identified DARs (Table S4), with mature NK cells exhibiting the highest number of upregulated DARs (Figure S8C). GO enrichment analysis of these upregulated DARs revealed that pathways related to “immune defense and killing,” “immune regulation,” and “immune cell mobilization” were significantly enriched in mature NK cells, whereas inflammation and stress-related pathways were downregulated (Figure 5D). To deeply investigate the transcriptional regulatory basis, we performed motif deviation analysis based on scATAC-seq data (Figure 5E). Results showed that in EX, motif accessibility was significantly enhanced for TFs closely related to NK cell immune function, including key regulators driving cytotoxic differentiation such as *RUNX3*, *EOMES*, and *TBX21* (T-bet), as well as *ETS1*, which is involved in innate lymphoid cell development and activation(*56*). Conversely, motif accessibility for inflammation and stress-related TFs (*RELA*, *RELB*, *NFKB1*, *JUND*, *FOS*, *FOSL2 and BACH1*)(*56*) was generally reduced. Footprint analysis confirmed the decreased accessibility of *RELA* and *RELB* transcription factor binding sites (Figure S8D). Notably, in mature NK cells, chromatin accessibility was significantly enhanced in EX at the intronic regulatory region of *STAT1*, a core mediator of type I interferon signaling, and the promoter region of *CXCR4*, a known exercise-responsive factor(*57*) (Figure 5F). Furthermore, accessibility was significantly reduced at the promoter regions of *RELB*, associated with non-canonical NF-κB pathway activation, and *CCL5*, an important inflammatory chemokine (Figure S8E). In summary, these results demonstrate that regular exercise reshapes NK cells into a more robust effector state, characterized by enhanced cytotoxic and mobilization capacities alongside reduced levels of inflammation and stress.

To investigate whether common regulatory mechanisms exist for the cytotoxic functions of adaptive and innate immunity under exercise, we compared transcription factor activities between CD8^+^ CTL (adaptive immunity) and mature_NK (innate immunity) cells (Figure 5G). Venn analysis revealed that both cell populations shared a core set of significantly upregulated transcription factors, including *RUNX3*, *EOMES*, *TBX21*, *FOXO3*, and *ETS2*. This provides a unified transcriptional regulatory basis for the synergistic enhancement of cytotoxic functions across different immune branches by regular exercise.

### Regular Exercise Potentiates B Cell Antigen Presentation via Epigenetic Reprogramming

Given the positive impact of regular exercise on vaccination responses and antibody levels(*58*, *59*), we employed single-cell multi-omics technology to conduct an in-depth analysis of B cells, the core executors of humoral immunity. B cells were annotated into eight subtypes (Figures S9A, S4E, and S4J): switched memory B cells (Switched_Bm), naive B cells (Naive_B), pre-switched memory B cells (pre-Switched_Bm), unswitched memory B cells (Unswitched_Bm), atypical memory B cells (Atypical_Bm), transitional B cells (Transitional_B), plasma cells, and plasmablasts. Cell proportions did not differ significantly between EX and SED across these subsets (Figure S9B).

Next, we performed DEG analysis and GO enrichment analysis (Table S3). Upregulation of pathways related to antigen processing and presentation, humoral immune response, and chemotaxis was observed in naive_B, switched_Bm, and transitional_B (Figure S9C). Notably, MHC molecules were significantly upregulated in EX across multiple B cell subsets: *HLA-DQB1* in naive_B, switched_Bm, and unswitched_Bm; *HLA-DQA1* in transitional_B; and *HLA-F* in unswitched_Bm (Figure S9D). Additionally, the chemokine *CCL5* and *S100A9* were significantly upregulated in switched_Bm cells, and the actin regulator *COTL1* was upregulated in transitional_B cells (Figure S9D). To resolve the epigenetic regulatory basis, we performed DAR analysis (Table S4). Results showed that changes in chromatin accessibility were highly concentrated in naive_B cells and switched_Bm (Figure S9E). GO enrichment analysis of the upregulated DARs in these two subsets revealed that both were enriched in pathways related to antigen presentation and positive regulation of B cell activation (Figure S9F).

To dissect the transcriptional regulation potentially influenced by these epigenetic alterations, we analyzed TF activity in naive_B and switched_Bm. In naive_B, we observed significant upregulation in the activity of key regulators. This included *BCL11A*(*60*), a critical determinant of B-cell lineage fate; *RUNX1*(*61*), essential for lymphocyte development; and the core master regulators of early B-cell commitment, *SPI1* (PU.1)(*62*) and *EBF1*(*63*) (Figure S9G). switched_Bm showed upregulation of NF-κB family members (*RELA*, *RFLB* and *NFKB1*), *IRF4*, as well as *SPI1* and *EBF1* (Figure S9H); NF-κB and *IRF4* are master regulators of memory B cell formation, class switching, and plasma cell differentiation(*64*, *65*). We focused our analysis on the regulatory regions of genes upregulated in the transcriptome. The results showed that in EX, chromatin accessibility at the promoter region of the key antigen presentation gene *HLA-DQB1* was significantly enhanced in both naive_B (Figure S9I) and switched_Bm (Figure S9J). Additionally, in switched_Bm, promoter accessibility of the chemokine gene *CCL5* also showed a synchronous increase (Figure S9K).

Overall, this study demonstrates that although regular exercise does not markedly alter the proportions of B cell subsets, it may enhance their antigen presentation and immune activation capabilities through chromatin remodeling and changes in gene expression. Recent research has highlighted the critical role of B cell antigen presentation in effective immunization and antiviral defense(*66*). Therefore, our findings suggest that the robust immune response observed in exercised individuals could be attributed to exercise-induced chromatin reprogramming and transcriptional modifications in B cells.

### Altered Intercellular Communication

Given the significant enhancement of antigen-presenting cells (APCs) function in EX, particularly in cMono, cDC, and B cells, we employed the CellChat algorithm to analyze whether this change remodeled the immune cell-cell interaction network at the systems level(*67*). Our analysis revealed that myeloid cells mediated the highest number of intercellular interactions among all cell types. Moreover, the most pronounced differential interaction strength between EX and SED was observed for myeloid cell–CD8^+^ T cell pairs (Figures 6A and 6B). At the pathway level, we observed a clear pattern: information flows related to inflammation and immune suppression (THBS, RESISTIN and IL6) were dominant in SED, whereas signals driving chemotaxis, antigen presentation, and cell adhesion (CCL, MHC-II and PECAM1) were markedly enhanced in EX (Figure 6C). Human resistin is a key regulator directing inflammatory responses in various immune cells(*68*). Cell communication network showed that myeloid cells were the core signal source for this pathway, and their signal output was significantly lower in EX. (Figure S10A). The interaction intensity of key ligand-receptor axes in this pathway, RETN–CAP1 and RETN–TLR4, was weakened in EX. The downregulation of *RETN* gene expression was also observed at the transcript level in myeloid cells (Figures S10B and S10C).

**Figure 6.**
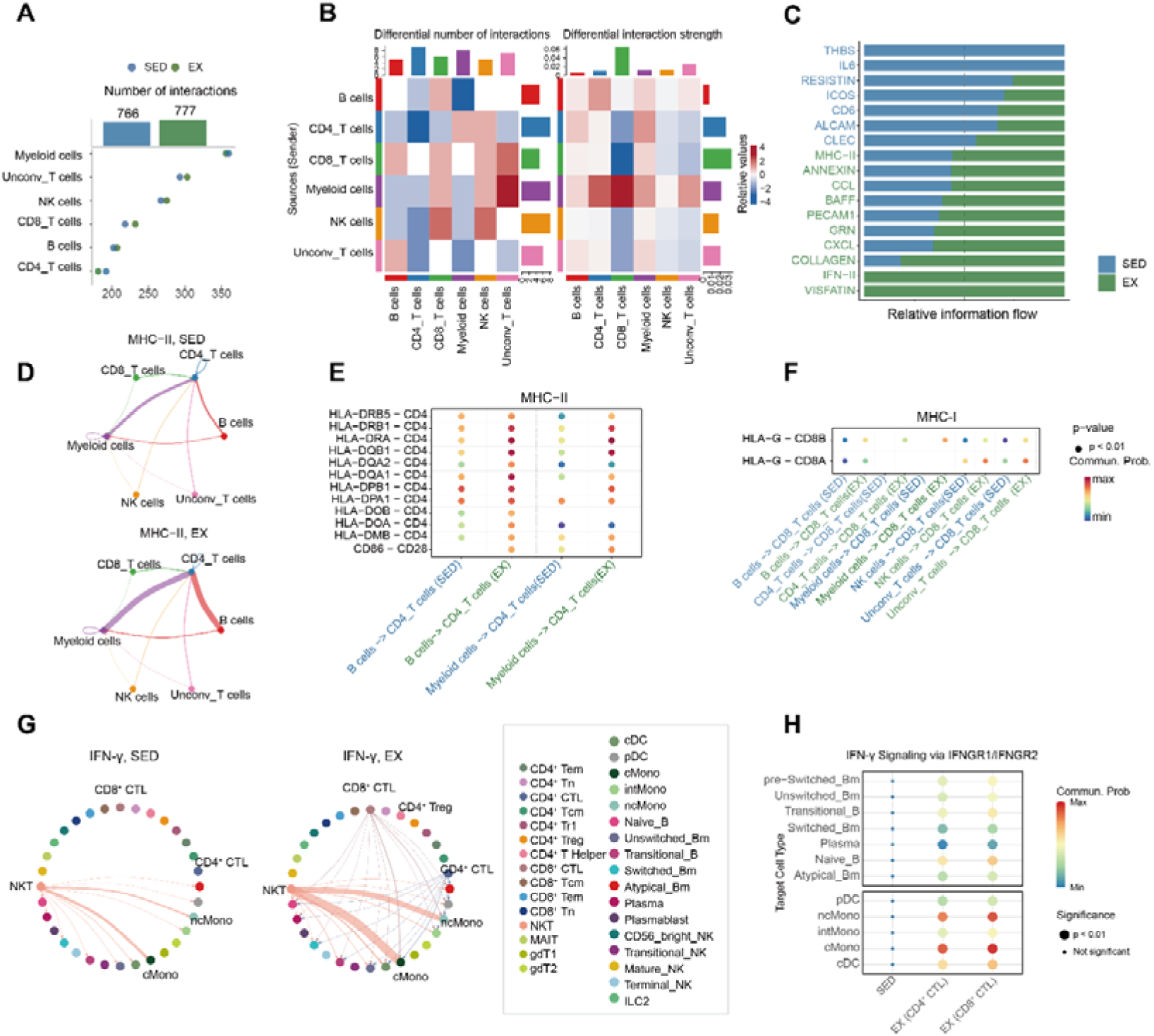
Remodeling of immune cell communication networks associated with regular physical activity. (A) Bar plot showing the total number of intercellular interactions in SED and EX, with dot plots indicating the number of interactions mediated by each major immune cell type. (B) Heatmaps showing the differential number of interactions (left) and differential interaction strength (right) between major immune cell types (EX vs. SED). Red indicates increased and blue indicates decreased interactions in EX. (C) Bar plot of relative information flow for each signaling pathway in SED and EX. Green and blue bars represent EX and SED, respectively. (D) Cell-cell communication network plots showing MHC-II signaling among major immune cell types in SED (top) and EX (bottom). Line thickness represents interaction strength. (E) Bubble plot showing communication probabilities of MHC-II ligand–receptor pairs between B cells and myeloid cells to CD4^+^ T cells in SED and EX. Dot size represents communication probability; color indicates statistical significance. (F) Bubble plot showing communication probabilities of MHC-I ligand–receptor pairs (HLA-G–CD8A and HLA-G–CD8B) from multiple cell types to CD8^+^ T cells in SED and EX. Dot size represents communication probability; color indicates statistical significance. (G) Cell-cell communication network plots showing IFN-γ signaling among immune cell subsets in SED (left) and EX (right). Line thickness represents interaction strength. (H) Bubble plot showing communication probabilities of IFN-γ signaling via IFNGR1/IFNGR2 from CD4^+^ CTL and CD8^+^ CTL to myeloid and B cell subsets in SED and EX. Dot size represents communication probability; color indicates statistical significance.

Exercise significantly enhanced intercellular signaling associated with antigen presentation. Compared to SED, EX exhibited a generalized enhancement of MHC-II signaling from B cells and myeloid cells to CD4^+^ T cells (Figures 6D and 6E). Subset analysis further pinpointed that interactions involving cMono, naive_B cells, and switched_Bm cells contributed most prominently within this pathway (Figure S10D). Besides, MHC-I signaling from various cell subsets (B cells, CD4^+^ T cells, myeloid cells, NK cells, and unconv_T cells) to CD8^+^ T cells showed a global increase.The enhancement of signaling mediated by the HLA-G–CD8A/B pair was the most pronounced (Figure 6F), primarily driven by interactions between cMono and CD8^+^ Tn cells (Figure S10E). Notably, exercise also potentiated the signaling network mediated by IFN-γ. In EX, IFN-γ signaling from NKT cells to multiple myeloid and B cell subsets was markedly enhanced (Figure 6G). In parallel, cytotoxic T cells (CD4^+^ and CD8^+^ CTLs) also strengthened their signaling via the IFNG–receptor axis to these same innate immune cell populations (Figure 6H). IFN-γ, a pivotal cytokine that bridges innate and adaptive immunity by driving CD4^+^ T cell differentiation toward the Th1 lineage, enhancing the cytotoxicity of CD8^+^ T cells, and serving as a core effector molecule for NK cell function(*69*, *70*).

Notably, exercise also potentiated signaling networks associated with tissue homeostasis and immune microenvironment organization. The COLLAGEN signaling pathway was markedly enhanced in EX, predominantly driven by unconv_T cells(Figures S10F). This was evidenced by the coordinated upregulation of both the key ligand-receptor pair COL6A2–CD44 and its corresponding ligand gene expression (Figures S10F-H). Collagen-mediated signaling is a well-established regulator of cell adhesion, migration, and tissue homeostasis(*71*). In addition, the CCL chemokine signaling pathway was significantly enhanced in EX, manifested primarily as increased signal output from other major immune cell lineages to myeloid cells (Figures S10I and S10J). CCL family chemokines are central orchestrators of immune cell recruitment and spatial positioning within tissues(*72*). In summary, regular exercise remodels the intercellular communication network among immune cells. MHC signaling from APCs to T cells was enhanced, accompanied by increased IFN-γ signaling from NKT and cytotoxic T cells to myeloid and B cell subsets, promoting T cell activation and effector function. Pathways involved in immune recruitment (CCL) and tissue organization (COLLAGEN) were also upregulated, while pro-inflammatory signals such as RESISTIN and IL6 were attenuated. These findings indicate that exercise coordinately enhances antigen presentation and T cell-mediated immunity while reducing chronic inflammation.

## Discussion

Physical inactivity is a risk factor for metabolic disorders, immune dysfunction, and all-cause mortality(*1*, *73*). In contrast, regular physical activity is a well-established, effective intervention to mitigate these risks(*74*). However, the underlying cellular and molecular mechanisms that confer protection in free-living populations remain incompletely characterized. Our study reveals that regular exercise not only significantly enhances systemic fatty acid oxidation efficiency and endogenous antioxidant defense, but also induces widespread epigenetic remodeling at the single-cell level, selectively optimizing antigen presentation and cytotoxic functions. Importantly, intercellular communication undergoes a systemic shift. This transition is characterized by the downregulation of pro-inflammatory mediators, coupled with the amplification of homeostatic pathways, particularly those related to MHC-mediated antigen presentation and IFN-γ signaling. Collectively, these findings reveal how regular exercise reshapes the internal milieu across the dimensions of metabolic homeostasis and immune function, providing a novel molecular perspective on its systemic health benefits.

EX exhibit a distinct metabolic signature characterized by elevated FAO intermediates, such as O−Adipoylcarnitine and ketone bodies, alongside a reduction in TAG levels. These findings indicate a significant enhancement in systemic lipid turnover and utilization efficiency. This metabolic phenotype aligns closely with the clinical improvements in lipid profiles (decreased TG and increased HDL-C), collectively forming the physiological basis for exercise-induced cardiometabolic protection(*75*). Furthermore, we observed an increase in endogenous antioxidants, including betaine and 3-hydroxyanthranilic acid, reflecting an augmented buffering capacity against oxidative stress in response to elevated metabolic flux(*36*). Notably, recent evidence has identified betaine as an endogenous TBK1 inhibitor that functions as an “exercise mimetic,” exerting anti-inflammatory and pro-longevity effects(*10*). The significantly elevated betaine levels observed in EX not only provide real-world human evidence for this mechanism but also suggest that betaine has the potential to serve as a biomarker of exercise adaptation and a promising therapeutic target for clinical intervention. Several metabolites upregulated in this study have been previously established as potent immunomodulatory signaling molecules or epigenetic regulators. For instance, β-hydroxybutyrate (BHB) functions as an endogenous histone deacetylase (HDAC) inhibitor that suppresses NLRP3 inflammasome activation in macrophages, thereby exerting systemic anti-inflammatory effects(*76*). Furthermore, GABA has been shown to directly regulate lymphocyte proliferation and inflammatory cytokine secretion through functional receptors expressed on the surface of immune cells(*77*). These established links suggest that exercise-induced systemic metabolic remodeling might contribute to the optimization of immune cell functions. Although the direct causal relationship between these metabolites and immune cells was not explicitly tested in this study, these observations provide a robust foundation for hypothesis generation in future mechanistic research.

Regular exercise not only reshapes systemic metabolic homeostasis but also orchestrates a functional reinforcement of the professional APCs landscape. Within the myeloid compartment, monocytes and DCs exhibit elevated expression of antigen-presentation-related genes. In cMono specifically, we observed enhanced chromatin accessibility at *HLA-DQB1* and *HLA-DQA1* loci, synchronized with the increased regulatory activities of master transcription factors governing antigen presentation and vascular homeostasis, such as *SPI1*, *IRF1*,and *KLF4*. This enhanced antigen presentation capacity enables more efficient antigen capture and presentation, facilitating the priming of adaptive immune responses and maintaining immune cells in a heightened state of alertness. Beyond the myeloid compartment, such epigenetic remodeling extends to adaptive immune subsets, where naive_B and switched_Bm cells also demonstrate coordinated increases in *HLA-DQB1* expression and regulatory element accessibility. Such cross-lineage remodeling aligns with our cell-cell communication analysis, which reveals a significant amplification of MHC-I and MHC-II signaling from APCs to T cells in EX. Given that antigen-presentation efficiency serves as a critical rate-limiting step in adaptive immune responses(*78*, *79*), the convergent enhancement of antigen presentation capacity across cMono and B cell subsets, coupled with the amplified APCs-T cell crosstalk, may collectively underlie the enhanced immune surveillance and reduced susceptibility to infection documented in physically active populations(*5*, *58*).

In T cell subsets, regular exercise is manifested as a distinct transition in effector function. For instance, in CD8^+^ CTLs, increased activities of master transcription factors driving effector programs, such as *TBX21* and *EOMES*, are observed, accompanied by the up-regulation of downstream target genes (*GZMB*) and increased chromatin accessibility at their regulatory elements. These alterations occur in synchronization with the systemic enhancement of TCR and IFN-γ response pathways, suggesting that regular exercise may enhance the sensitivity of effector T cells to activation signals by optimizing the epigenetic landscape. Furthermore, the increased motif accessibility of factors such as *TCF7* in CD8^+^ Tn cells suggests that, while reinforcing the effector reservoir, exercise may also contribute to maintaining or optimizing the stemness and long-term adaptive potential of the naive T cell pool(*80*, *81*). Similar regulatory features, such as the increased openness of *EOMES*/*TBX21* motifs, are also reflected in mature NK cells.

Cell communication analysis indicates that regular exercise can systematically remodel the intercellular interaction network, thereby coordinating metabolic regulation and immune remodeling. This study found that EX simultaneously enhanced immune surveillance and coordination pathways while suppressing chronic inflammatory signaling. Specifically, EX upregulated key immune-coordination pathways such as interferon signaling, antigen presentation (MHC-I/II), and chemokine activity, which may contribute to improved immune defense and tissue communication. Concurrently, within this network of cellular interactions, the pro-inflammatory network centered on resistin signaling and dominated by myeloid cells was inhibited. Research has established that resistin acts not only as a potent pro-inflammatory factor, upregulating inflammatory mediators such as tumor necrosis factor-α (TNF-α) and IL-6(*82*), but also directly targets the vascular wall, inducing endothelial dysfunction and promoting smooth muscle cell proliferation, thus accelerating the progression of pathological processes including atherosclerosis and vascular restenosis(*83*, *84*). Therefore, this study suggests that regular exercise may provide a protective regulatory foundation for the cardiovascular system at the level of cellular interactions, potentially through the inhibition of this key pro-inflammatory network.

In summary, this study employs multi-omics to describe exercise-related changes in circulating metabolites and lipids, alongside cell-type-resolved transcriptional and epigenetic landscapes of immune cells. These are characterized by enhanced lipid oxidation and antioxidant metabolic capacity, alongside functionally optimized immune states guided by epigenetic reprogramming. We provide multidimensional map of spontaneous exercise-induced molecular adaptations across metabolic, epigenetic, and immune dimensions, thereby deepening our understanding of exercise-mediated health promotion mechanisms and identifying potential biomarkers and regulatory pathways for chronic disease prevention and management.

## Limitations

Several limitations of this study warrant consideration. First, the assessment of physical activity levels relied on self-reporting; although this is a standard methodology in large-scale epidemiological research, it remains susceptible to recall and social desirability biases. Second, the study population consisted primarily of young, healthy individuals. While this focus facilitates the exclusion of confounding factors and reveals the core adaptive mechanisms of regular exercise, caution should be exercised when generalizing these conclusions to elderly, obese, or clinical populations. Third, although the single-cell multi-omics sample size is substantial relative to comparable studies and all analyses were statistically adjusted for sex and BMI, the higher proportion of females in the sedentary group may still introduce residual confounding. Expanding the sample size in future cohorts will aid in achieving better baseline balance and more granular subgroup analyses. Nevertheless, our study provides a multi-omics characterization of molecular adaptations to regular exercise in a natural population cohort, laying a foundation for future mechanistic and translational studies.

## Methods

### Data Source

All data for this study were derived from the CIMA project, a publicly available cohort that enrolled 428 adults with systematic multi-dimensional data collection(*21*). Our analysis utilized the following data modalities: (1) CIMA sample information metadata and physical activity questionnaires, (2) clinical blood biochemical markers, (3) targeted plasma lipidomics and metabolomics profiles, and (4) raw sequencing data matrices from paired scRNA-seq and scATAC-seq performed on PBMCs.

### Assessment of Physical Activity and Grouping

Physical activity was assessed across three dimensions (intensity, frequency, and duration) using a self-reported questionnaire within the CIMA cohort. Exercise intensity was determined by the item, “What intensity of exercise do you usually engage in?” (options: low, moderate, high, or almost none). Frequency (times/week) and session duration were derived from the items “Exercise habit” and “What is the average daily exercise duration?”, respectively. To obtain a conservative estimate, the lower bounds of the reported intervals were used for calculation (e.g., “3-4 times/week” was counted as 3; “1-2 hours” was taken as 60 minutes). Based on the WHO Guidelines on Physical Activity and Sedentary Behaviour(*23*), participants who engaged in exercise ≥ 3 times per week and met the recommended exercise volume were classified into the regular exercise group (EX, n = 40), and those reporting no exercise and a sedentary lifestyle were classified into the sedentary control group (SED, n = 46).

### Differential Analysis of Plasma Metabolites and Lipids

The raw targeted metabolomics and lipidomics data matrices were preprocessed and normalized using the MetaboAnalyst 6.0 online platform(*85*). Metabolites or lipid species missing in more than one-third of samples were removed. Remaining missing values were imputed using the minimum value detected for each molecule. The data were then subjected to Auto-Scaling (unit variance scaling), i.e., each molecule was mean-centered and divided by its standard deviation, to eliminate scale differences and give equal analytical weight to all molecules.To identify molecules associated with exercise habits, a combined screening strategy was applied. First, linear models were fitted using the limma package in R, adjusting for sex and BMI, and molecules with *p* < 0.05 and |log_2_FC| ≥ 0.15 between exercise groups were selected. Partial least squares discriminant analysis (PLS-DA) was performed using the MetaboAnalystR package (version 4.0) to calculate variable importance in projection (VIP) scores(*86*). Ultimately, metabolites and lipids that satisfied both criteria—VIP scores ≥ 1 and the linear-model thresholds—were defined as exercise-associated differential molecules (Supplementary Table S2).

### Pathway Analysis of Plasma Metabolites

To uncover the potential biological processes underlying the differential metabolites, Kyoto Encyclopedia of Genes and Genomes (KEGG) pathway enrichment analysis was conducted for the exercise-associated differential metabolites using the MetaboAnalyst 6.0 platform(*85*). Enrichment significance was evaluated with the hypergeometric test, and p-values were corrected for multiple testing using the false discovery rate (FDR). To identify key metabolic alterations, we focused on pathways that passed a significance threshold of FDR-corrected p-value < 0.05 in our enrichment analysis.

### Single-cell RNA-seq data processing

This study used processed single-cell gene expression count matrices. All analyses were performed using Scanpy (version 1.10.3)(*87*). Quality control retained cells with 500–6000 genes, 1000–25,000 unique molecular identifiers (UMIs), and mitochondrial content < 10% (Figure S1A). Doublets were identified and removed using Scrublet (expected doublet rate = 0.06)(*88*). After filtering, data were normalized to 10,000 counts per cell, log-transformed, and scaled. The top 3,000 highly variable genes were selected for principal component analysis, and batch effects were corrected using Harmony(*89*).

### Unsupervised clustering and subclustering

After batch-effect correction, unsupervised clustering was performed using the Louvain algorithm(*90*) (resolution = 0.75), and visualized with UMAP. Based on canonical marker genes, six major cell populations were identified: CD4^+^ T, CD8^+^ T, unconventional T, B, myeloid, and NK cells. Each major population was then extracted and re-clustered separately with subset-optimized resolutions. Subcluster identities were assigned based on differential expression and known subtype-specific markers. In total, 34 distinct cell subpopulations were identified across the dataset.

### Differential abundance analysis using scRNA-seq data

To evaluate exercise-associated changes in immune cell composition, differential abundance analysis was performed using Milo(*32*). A k-nearest neighbor graph (k = 30) was constructed from the first 50 principal components, and cells were partitioned into overlapping neighborhoods (makeNhoods, prop = 0.2). Differential abundance testing between EX and SED was performed using a negative binomial GLM, with TMM normalization and adjustment for sex and BMI. Significant neighborhoods were defined as those with FDR < 0.05, and their log_2_FC values were used for downstream interpretation.

### Differential Expression Analysis of Cell Subpopulations

For the comparison of gene expression differences between the EX and SED groups across cell subpopulations, we employed the linear model-based limma package for statistical testing(*91*). For each subpopulation, we constructed a linear model in which gene expression was the dependent variable, exercise group was the primary predictor, and sex and BMI were included as covariates. From the fitted model, the log_2_FC and raw p-value were extracted for every gene. To control for false positives, we adopted the Benjamini-Hochberg (BH) procedure for multiple testing correction of the p-values. In our analysis, a gene was considered a significant DEG within a given subpopulation if it showed |log_2_FC| ≥ 0.25 together with an adjusted p-value < 0.05.

### Module scoring using scRNA-seq data

To evaluate activity differences in specific biological processes across immune cell subpopulations, Scoring cells for specific gene modules was based on the Scanpy sc.tl.score_genes method, which computes the difference between the average expression of target genes and that of a matched random background per cell. For our analysis, we sourced gene sets from the Molecular Signatures Database (MSigDB)(*92*).To assess statistical differences in module scores between the EX and SED groups, we first computed the mean module score per subpopulation. We then fitted linear models comparing groups, adjusting for sex and BMI.

### Gene regulatory network analysis

Transcriptional regulatory networks in immune cell subsets were inferred using pySCENIC (v0.12.1)(*93*). GRNBoost was used to identify TF-target gene co-expression modules, which were then refined against the cisTarget database to retain only genes with relevant TF-binding motifs(*94*). Regulon activity was quantified by areas under the curve per cell (AUCell), and differential regulon activity between EX and SED was assessed with limma (adjusted p < 0.05, |log_2_FC| ≥ 0.25).

### scATAC-seq quality control

We used ArchR (version 1.0.2) to analyze single-cell ATAC-seq data(*95*). First, Arrow files were generated for each sample from the raw sequencing fragments following the default settings. Subsequently, stringent cell-level quality control was applied. Our cell quality control was based on the parameters below: TSS enrichment ≥ 10, number of unique fragments between 1×10^3^ and 1×10^5^, nucleosome signal < 4, and blacklist ratio < 0.05. Doublets were identified and removed using ArchR’s default parameters, resulting in a high-quality chromatin accessibility matrix for downstream analysis.

### scATAC-seq dimensionality reduction and clustering

To construct a single-cell chromatin accessibility landscape, we first performed dimensionality reduction on the cell-by-peak matrix using latent semantic indexing (LSI) to capture dominant patterns of chromatin accessibility variation. To mitigate technical variability introduced during sample preparation, we applied the Harmony(*89*) algorithm to integrate the reduced-dimension embeddings across samples.Using the integrated low-dimensional representation, we partitioned cells into communities via shared nearest neighbor (SNN)-based graph clustering. An initial clustering step was performed at a uniform resolution of 0.8 to delineate major cell categories. To further resolve cellular heterogeneity and enhance the detection of small-scale cell states, we carried out sub-clustering on selected populations derived from the initial clustering, with resolutions adapted between 0.3 and 1.5 according to the intrinsic structure of each cell group. As the final step in our single-cell ATAC-seq analysis, all cells were projected into a two-dimensional space for visualization using uniform manifold approximation and projection (UMAP), providing an intuitive representation of cell-state distributions and community relationships.

### scATAC–seq gene activity scores

Gene activity was quantified with ArchR(v1.0.2), which computes scores by correlating accessibility across gene-linked regions. Following this, the MAGIC algorithm was applied to smooth the scores and mitigate noise inherent in sparse scATAC-seq data(*96*).

### Integrating scATAC-seq and scRNA-seq Data to Infer Cell Identity

To accurately annotate cell types in the scATAC-seq data, we performed an integrative analysis against a matched scRNA-seq reference. This was achieved using the addGeneIntegrationMatrixfunction in ArchR. This process was conducted in two sequential steps: first, an unconstrained integration was carried out to broadly map each scATAC-seq cell cluster to the major cell types defined in the scRNA-seq data. Subsequently, based on this initial mapping, A second, more constrained integration was then carried out by re-executing addGeneIntegrationMatrix with updated broad group labels. This refinement enabled finer multimodal alignment and produced final annotations at high resolution, ensuring consistency with the transcriptomic profiles.To independently assess the reliability of the annotations, we employed the Gene Activity Score (GAS) matrix derived from the scATAC-seq data for validation. By comparing the distribution of annotated cells in the gene-activity space with that of the corresponding scRNA-seq reference cells in gene-expression space, we further confirmed the accuracy of the cross-modal annotation.

### scATAC–seq pseudobulk replicate generation and peak calling

To support differential analysis across cell clusters and conditions, we first constructed pseudobulk samples. This involved running the addGroupCoverages function with tailored parameters to create non-overlapping replicates for each cell group. These replicates were then consolidated via addReproduciblePeakSet to generate a consensus peak matrix. For precise peak identification, we used MACS2(*97*). The resulting unified peak set provided the foundation for all subsequent differential accessibility testing.

### scATAC–seq genomic regions annotation

For the identified chromatin-accessible regions (peaks), We annotated the called peaks by applying the annotatePeak function from the ChIPseeker package package with default parameters(*98*), in order to determine the nearest associated genes and genomic features for each peak.

### scATAC motif enrichment and motif deviation analysis

We carried out transcription factor motif enrichment followed by chromatin accessibility deviation analysis using the unified pseudobulk peak set. For motif annotation, we utilized the CIS-BP motif database (via ChromVAR)(*99*), JASPAR2020(*100*), and HOMER(*101*). Additionally, motif accessibility deviation scores were calculated for each cell using the chromVAR algorithm implemented in ArchR, enabling systematic assessment of changes in transcription factor binding activity across different cellular states.

### Identification of differential peaks

To detect differentially accessible chromatin regions between groups, we employed two complementary approaches. First, we used the getMarkerFeatures function with parameters useMatrix = “PeakMatrix”, bias covariates including “TSSEnrichment” and “log10(nFrags)”, and testMethod = “wilcoxon” to identify differential peaks between groups, applying thresholds of FDR ≤ 0.1 and log_2_FC ≥ 0.25. Second, for differential transcription factor activity analysis, we applied the same function with useMatrix = “MotifMatrix” and thresholds of FDR ≤ 0.01 and MeanDiff ≥ 0.05.

### Gene Ontology Enrichment Analysis of differential peaks

To interpret the potential biological functions of differential chromatin accessible regions, we performed GO enrichment analysis using the R package rGREAT(*102*). This package associates genomic regions with nearby genes and evaluates their enrichment significance across Biological Process, Molecular Function, and Cellular Component terms using a hypergeometric test. The analysis was conducted with the human genome as background, applying default parameters, We selected terms passing the significance threshold of an FDR-adjusted p-value < 0.05 as significantly enriched.

### scATAC–seq TF Foot-print analysis

We performed transcription factor footprinting analysis based on Tn5 transposase cleavage profiles across genomic motif regions. First, the sequence bias of Tn5 was subtracted from the raw cleavage signal to obtain a corrected footprint profile. Motif positions were determined using a combination of the CIS-BP database(*103*) (from chromVAR human_pwms_v1), and the JASPAR2020 database. Subsequently, footprint signals were normalized using the average insertion density within ± 200–250Lbp regions centered on each motif. The normalized results were aggregated by pseudo-replicate, and their means and standard deviations were calculated and visualized. Finally, to assess differences in footprint signals between groups, We chose the nonparametric Wilcoxon rank-sum test for comparisons between two groups, setting the significance level at *p* < 0.05.

### scATAC-seq Peak to gene linkage analysis

To identify peak-to-gene links, we performed prediction analysis using the addPeak2GeneLinks function in ArchR. The analysis was based on the batch-corrected low-dimensional embedding (specified via the reducedDims parameter) and computed the correlation between chromatin accessibility and gene expression. A correlation threshold of corCutOff = 0.4 was applied to screen for high-confidence linkages. The resulting significant associations were output as a GRanges object for subsequent visualization and functional interpretation.

### Cell-cell interaction analysis

We performed a systematic analysis of potential signaling communication among immune cell subpopulations using the CellChat R package(*67*). Communication probabilities between cell types were computed using default parameters. Signaling pathways were summarized and the global communication network was constructed using the standard CellChat workflow. To compare communication patterns between EX and SED, differential interaction strength was assessed with the rankNet function.

### Statistical Analysis

Detailed statistical methods for each analysis are provided in the corresponding Method subsections and figure legends. All statistical analyses were performed in R (v4.1.3) and Python (v3.9). Unless otherwise specified, statistical significance was defined as FDR < 0.05. In the figures, significance levels are denoted as \**p* < 0.05, \*\**p* < 0.01, \*\*\**p* < 0.001, \*\*\*\**p* < 0.0001.

## Acknowledgements

We thank all of our team members.

## Author contributions

J.L., X.S., S.G., J.Y., P.Y., and C.L. conceived the idea; C.L., P.Y., J.Y., Y.Z., L.L., P.G., and J.Z. supervised the work; X.S., J.L., S.G., S.X., Y.W., Y.Z., and W.Z. analyzed the data and performed visualization; X.S., J.L., S.G., P.Y., J.Y., Z.C., and C.L. wrote the manuscript.

## Data availability

The scRNA-seq and scATAC-seq data, plasma metabolomics and lipidomics data, and blood biochemistry data presented in this study were sourced from the CIMA cohort, hosted on the TrueBlood database within the CNGBdb portal (https://db.cngb.org/trueblood/cima/).

## Code availability

The code can be obtained from the corresponding author upon reasonable request. Information on publicly available software used is provided in the Methods.

