## Supplementary Figures for "Multi-omics Profiling Identifies Molecular and Cellular Signatures of Regular Physical Activity in Human Peripheral Blood"

#### SUPPLEMENTAL FIGURE LEGENDS

**Figure S1. Baseline characteristics and analysis of blood biochemical indices of the study population, related to Figure 1.**

(A) A table summarizing the baseline characteristics of participants (n=86) stratified by physical activity pattern.

(B) Beeswarm and box plots showing serum concentrations of glucose (GLU), high-density lipoprotein cholesterol (HDL-C), and triglycerides (TG) in SED and EX. Adjusted *p*-values are indicated.

(C) Beeswarm and box plots showing serum concentrations of creatinine (Cr), uric acid (UA), indirect bilirubin (IBIL), total bilirubin (Tbil), and direct bilirubin (DBIL) in SED and EX. Adjusted *p*-values are indicated.

**Figure S2**. **Quality control, batch effect evaluation, and annotation validation for single-cell multi-omics data, related to Figure 1.**

(A) Violin plots depicting key scRNA-seq quality control metrics between the SED (blue) and EX (green) cohorts.

(B) Violin plots of scATAC-seq quality control indicators comparing the SED and EX groups.

(C) UMAP plots displaying cell distribution in scRNA-seq (top) and scATAC-seq (bottom) datasets, with cells colored by group.

(D) UMAP plots showing the expression levels of key marker genes in scRNA-seq data.

(E) UMAP plots showing chromatin accessibility at the promoter regions of key marker genes in scATAC-seq data.

(F) Heatmap of main marker genes based on gene activity scores from scATAC-seq data.

(G) Beeswarm and box plots showing the proportion of myeloid cell subsets in SED and EX based on scRNA-seq (top) and scATAC-seq (bottom) data.

**Figure S3. Plasma lipid species profiles associated with physical activity patterns, related to Figure 2.**

(A) Violin plots overlaid with box plots showing the normalized intensity of membrane phospholipid subclasses (LPE, PC, PG, and PI) in SED and EX. *p*-values are indicated.

(B) Normalized intensity of two differentially abundant PE species, PE 20:4-22:5 and PE 20:4-22:1, in SED and EX. *p*-values are indicated.

**Figure S4. Marker gene expression and chromatin accessibility-based gene activity scores across immune cell subsets, related to Figures 3–5.**

(A–E) Dot plots showing the expression of canonical marker genes for myeloid (A), $CD8^+$ and unconventional T (B), $CD4^+$ T (C), NK (D), and B cell (E) subsets from scRNA-seq data. Dot size represents the fraction of cells expressing the gene; color indicates average expression level.

(F–J) GeneScoreMatrix heatmaps showing chromatin accessibility-based gene activity scores of subset-specific marker genes for myeloid (F), $CD8^+$ and unconventional T (G), $CD4^+$ T (H), NK (I), and B cell (J) subsets from scATAC-seq data. Color represents column Z-score; representative marker genes are labeled.

**Figure S5. Myeloid cell subset characterization in SED and EX, related to Figure 3.**

(A) Beeswarm and box plots showing the proportion of Myeloid cell subsets in SED and EX groups based on scRNA-seq and scATAC-seq data.

(B) A heatmap showing normalized module scores of functional pathways across myeloid cell subsets between the SED and EX groups(*$p$ < 0.05,**$p$ < 0.01,***$p$ < 0.001).

(C) Pie chart showing the distribution of DARs across Myeloid cell subsets.

(D) Genome browser tracks of chromatin accessibility at the promoter DAR of *CXCL8* (left), distal DAR of *CCL2* (middle), and promoter DAR of *CCL7* (right) in cMono between SED and EX. Highlighted regions indicate differentially accessible regions with decreased accessibility in EX. $Log_2FC$ and FDR values are indicated.

(E) Violin plot overlaid with box plot showing the expression level of *CXCL8* in cMono between SED and EX. $Log_2FC$ and adjusted $p$-value are indicated. ***$p$ < 0.001.

**Figure S6. $CD8^+$ and unconventional T cell subset characterization in SED and EX, related to Figure 4.**

(A) Beeswarm and box plots showing the proportion of $CD8^+$ and unconventional T cell subsets in SED and EX groups based on scRNA-seq and scATAC-seq data.

(B) Heatmap showing the number of upregulated and downregulated DEGs across $CD8^+$ and unconventional T cell subsets (EX vs. SED).

(C) Box plots showing pathway activity scores for T cell-mediated immunity, T cell-mediated cytotoxicity, T cell activation, and type II interferon production across $CD8^+$ and unconventional T cell subsets in SED and EX. Adjusted p-values are indicated. *$p$ < 0.05, **$p$ < 0.01, ***$p$ < 0.001.

(D) Heatmap showing scaled expression levels of cytotoxic effector genes in $CD8^+$ CTL, $CD8^+$ Tem, and NKT cells between SED and EX. *$p$ < 0.05, **$p$ < 0.01, ***$p$ < 0.001.

(E–G) TF motif deviation analysis in $CD8^+$ CTL (E), MAIT (F), and $CD8^+$ Tn (G) cells. Each point represents a TF motif; x-axis shows mean chromatin accessibility difference (EX vs. SED); y-axis shows $–log_{10}$(FDR). Green and blue points indicate TF motifs with significantly increased and decreased accessibility in EX, respectively; gray points indicate non-significant motifs.

(H) Genome browser tracks of chromatin accessibility at the promoter DAR of *PRF1* in $CD8^+$ CTL cells between SED and EX.

(I) Genome browser tracks of chromatin accessibility at the intronic DAR of *TCF7* in $CD8^+$ Tn cells between SED and EX.

**Figure S7. $CD4^+$ T cell subset characterization in SED and EX, related to Figure 4.**

(A) UMAP visualization of $CD4^+$ T cell subsets identified by scRNA-seq (left) and scATAC-seq (right), colored by cell type.

(B) Box plots showing the proportion of $CD4^+$ T cell subsets in SED and EX based on scRNA-seq (top) and scATAC-seq (bottom) data. Adjusted *p*-values are indicated. ns, not significant. $*p < 0.05$.

(C) Beeswarmand box plots showing the distribution of $\log_2$FC differences in cell neighborhoods across $CD4^+$ T cell subsets (EX vs. SED), as determined by Milo differential abundance analysis. Box plots are created in a similar fashion as in Figure 1H.

(D) Heatmap showing the number of upregulated and downregulated DEGs across $CD4^+$ T cell subsets (EX vs. SED).

(E–F) Bar plots of GO enrichment analysis for upregulated DEGs in $CD4^+$ Tn (E) and $CD4^+$ T Helper (F) cells (EX vs. SED). Bar length represents $-\log_{10}$(adjusted *p*-value).

**Figure S8. NK cell subset characterization in SED and EX, related to Figure 5.**

(A) Box plots showing the proportion of NK cell subsets in SED and EX based on scRNA-seq (top) and scATAC-seq (bottom) data. Adjusted *p*-values are indicated. ns, not significant. $*p < 0.05$, $**p < 0.01$.

(B) Heatmap showing normalized pathway activity scores across NK cell subsets in SED and EX. Color represents normalized module score. $*p < 0.05$, $**p < 0.01$, $***p < 0.001$.

(C) Heatmap showing the number of DARs across NK cell subsets (EX vs. SED).

(D) TF footprint analysis showing Tn5 bias-subtracted normalized insertion profiles at

*RELA* (left) and *RELB* (right) motif centers in Mature_NK cells from SED and EX.

(E) Genome browser tracks of chromatin accessibility at the promoter DARs of *RELB* (left) and *CCL5* (right) in Mature_NK cells between SED and EX.

**Figure S9. B cell subset characterization in SED and EX, related to Figure 6.**

(A) UMAP visualization of B cell subsets identified by scRNA-seq (left) and scATAC-seq (right), colored by cell type.

(B) Beeswarmand box plots showing the distribution of $\log_2$FC differences in cell neighborhoods across B cell subsets (EX vs. SED), as determined by Milo differential abundance analysis.

(C) Bubble plot of GO enrichment analysis for DEGs across B cell subsets (EX vs. SED). Dot size represents $-\log_{10}$(adjusted *p*-value); color indicates regulation direction (pink: upregulated; blue: downregulated).

(D) Violin plots overlaid with box plots showing expression levels of selected genes across B cell subsets in SED and EX. $\text{Log}_2$FC adjusted *p*-values and statistical significance are indicated. $^*p < 0.05$, $^{***}p < 0.001$.

(E) Heatmap showing the number of DARs across B cell subsets (EX vs. SED).

(F) Bubble plot of GO enrichment analysis for upregulated DARs in Naive_B and Switched_Bm cells (EX vs. SED). Dot size represents gene ratio; color indicates $-\log_{10}$(FDR).

(G–H) TF motif enrichment analysis of upregulated peaks in Naive_B (G) and Switched_Bm (H) cells from EX. TFs are ranked by statistical significance. Violin plots showing motif deviation scores for selected TFs (*RELB*, *IRF4*, *SPI1*, and *EBF1*) in Switched_Bm cells between SED and EX are shown alongside (H). Adjusted p-values are indicated.

(I–K) Genome browser tracks of chromatin accessibility at the promoter DARs of *HLA-DQB1* in Naive_B (I) and Switched_Bm (J) cells, and *CCL5* in Switched_Bm (K) cells between SED and EX.

**Figure S10. Detailed intercellular communication networks in SED and EX,**

**related to Figure 6.**

(A) Cell-cell communication network plots showing RESISTIN signaling among major immune cell types in SED (left) and EX (right). Line thickness represents interaction strength.

(B) Bubble plot showing communication probabilities of RESISTIN ligand–receptor pairs (RETN–TLR4 and RETN–CAP1) from myeloid cells to all major immune cell types in SED and EX. Dot size represents communication probability; color indicates statistical significance.

(C) Violin plot overlaid with box plot showing the expression level of *RETN* in myeloid cells between SED and EX. $Log_2FC$ and adjusted p-value are indicated. ***$p < 0.001$.

(D–E) Cell-cell communication network plots showing MHC-II (D) and MHC-I (E) signaling at the single-cell subset level in SED (left) and EX (right). Line thickness represents interaction strength.

(F) Cell-cell communication network plots showing COLLAGEN signaling among major immune cell types in SED (left) and EX (right). Line thickness represents interaction strength.

(G) Bubble plot showing communication probabilities of the COL6A2–CD44 ligand–receptor pair from unconventional T cells to all major immune cell types in SED and EX. Dot size represents communication probability; color indicates statistical significance.

(H) Violin plot overlaid with box plot showing the expression level of *COL6A2* in unconventional T cells between SED and EX. $Log_2FC$ and adjusted p-value are indicated. ***$p < 0.001$.

(I) Cell-cell communication network plots showing CCL signaling among major immune cell types in SED (left) and EX (right). Line thickness represents interaction strength.

(J) Bubble plot showing communication probabilities of CCL ligand–receptor pairs (CCL5–CCR1 and CCL3–CCR1) from multiple immune cell types to myeloid cells in SED and EX. Dot size represents communication probability; color indicates statistical significance.

#### Figure S1

**A**

Participants' Characteristics at Baseline[a]

| Characteristic | Physical Activity Pattern, No. (%) of Participants: Total (n=86) | SED (n=46) | EX (n=40) | *p* value[b] |
|---|---|---|---|---|
| Age, y, weighted mean (95% CI) | 28.2 (27.2-29.2) | 28.9 (27.5-30.3) | 27.4 (26.0-28.8) | 0.14 |
| Sex, n (weighted %) | | | | < 0.05 |
| Male | 28 (32.6) | 9 (19.6) | 19 (47.5) | |
| Female | 58 (67.4) | 37 (80.4) | 21 (52.5) | |
| BMI, mean (SD) | 21.3 (±3.0) | 22.0 (±2.8) | 20.7 (±3.0) | < 0.05 |
| Lifestyle | | | | |
| Smoker, n (weighted %) | | | | 0.26 |
| Never | 82 (95.3) | 44 (95.7) | 38 (95.0) | |
| Past | 4 (4.7) | 2 (4.3) | 2 (5.0) | |
| Current drinker, n (weighted %) | | | | 0.15 |
| Yes | 16 (21.3) | 5 (16.2) | 11 (29.2) | |
| No | 70 (78.7) | 41 (83.8) | 29 (70.8) | |
| Sleep duration, n (weighted %) | | | | 0.56 |
| 6-8h | 72 (83.8) | 40 (87.0) | 32 (80.0) | |
| ≥ 8h | 14 (16.2) | 6 (13.0) | 8 (20.0) | |
| Dietary Habits, n (weighted %) | | | | 0.11 |
| Balanced Diet | 44 (51.1) | 19 (41.3) | 25 (62.5) | |
| Meat-Dominant Diet | 12 (14.0) | 9 (19.6) | 3 (7.5) | |
| Other | 30 (34.9) | 18 (39.1) | 12 (30.0) | |
| Medication Usage, n (weighted %) | | | | 0.08 |
| No Medication | 44(51.1) | 28 (60.9) | 16 (40.0) | |
| Used Medication | 44(48.9) | 18 (39.1) | 24 (60.0) | |

Abbreviations: BMI, body mass index(calculated as weight in kilogramsdivided by the height in meterssquared)

a Data were derived from the core cohort selected from the CIMA database, with a total of 86 participants included in this analysis.

b independent samples t-test within the same sex subgroup showed no significant difference in BMI between the two groups.

**B**

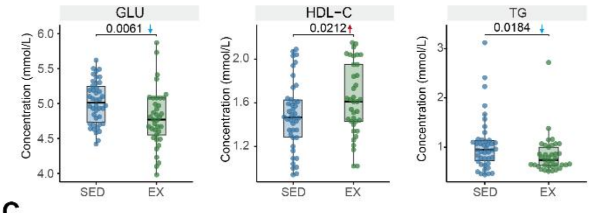


**C**

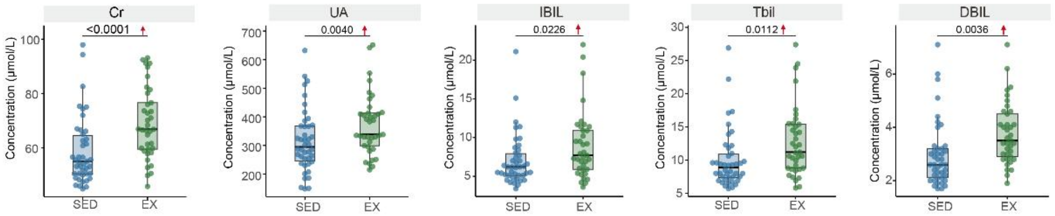

### Figure S2

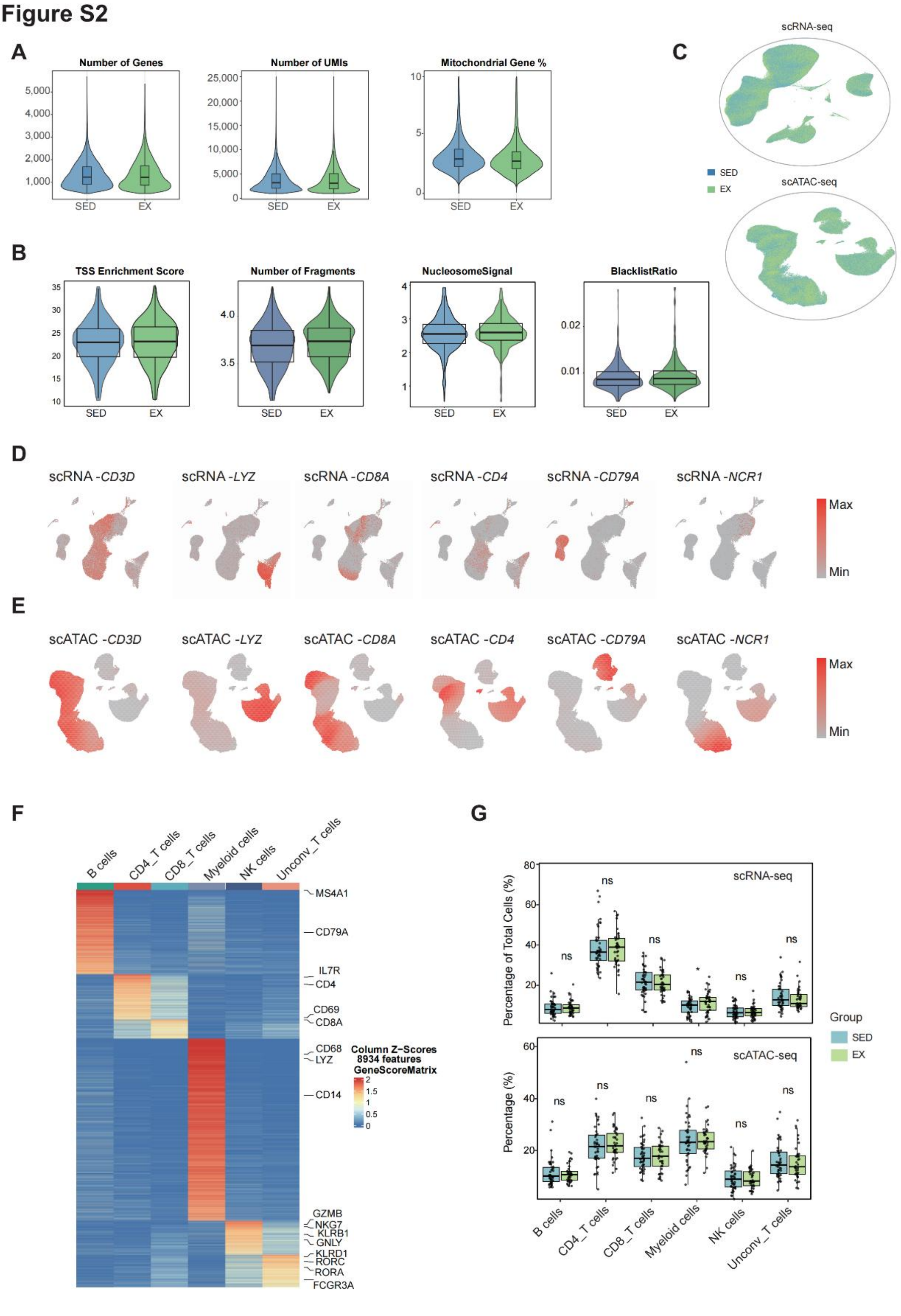

A
Number of Genes
Number of UMIs
Mitochondrial Gene %
SED
EX
B
TSS Enrichment Score
Number of Fragments
NucleosomeSignal
BlacklistRatio
C
scRNA-seq
scATAC-seq
D
scRNA -CD3D
scRNA -LYZ
scRNA -CD8A
scRNA -CD4
scRNA -CD79A
scRNA -NCR1
Max
Min
E
scATAC -CD3D
scATAC -LYZ
scATAC -CD8A
scATAC -CD4
scATAC -CD79A
scATAC -NCR1
F
B cells
CD4_T cells
CD8_T cells
Myeloid cells
NK cells
Unconv_T cells
MS4A1
CD79A
IL7R
CD4
CD69
CD8A
CD68
LYZ
CD14
GZMB
NKG7
KLRB1
GNLY
KLRD1
RORC
RORA
FCGR3A
Column Z-Scores
8934 features
GeneScoreMatrix
G
Percentage of Total Cells (%)
Percentage (%)
ns
Group

### Figure S3

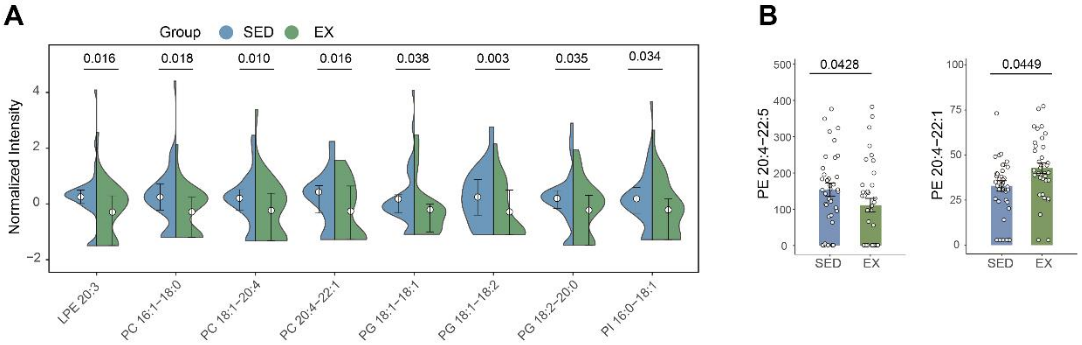
A
Group SED EX
0.016
0.018
0.010
0.016
0.038
0.003
0.035
0.034
Normalized Intensity
LPE 20:3
PC 16:1–18:0
PC 18:1–20:4
PC 20:4–22:1
PG 18:1–18:1
PG 18:1–18:2
PG 18:2–20:0
PI 16:0–18:1
B
0.0428
PE 20:4–22:5
SED
EX
0.0449
PE 20:4–22:1
SED
EX

### Figure S4

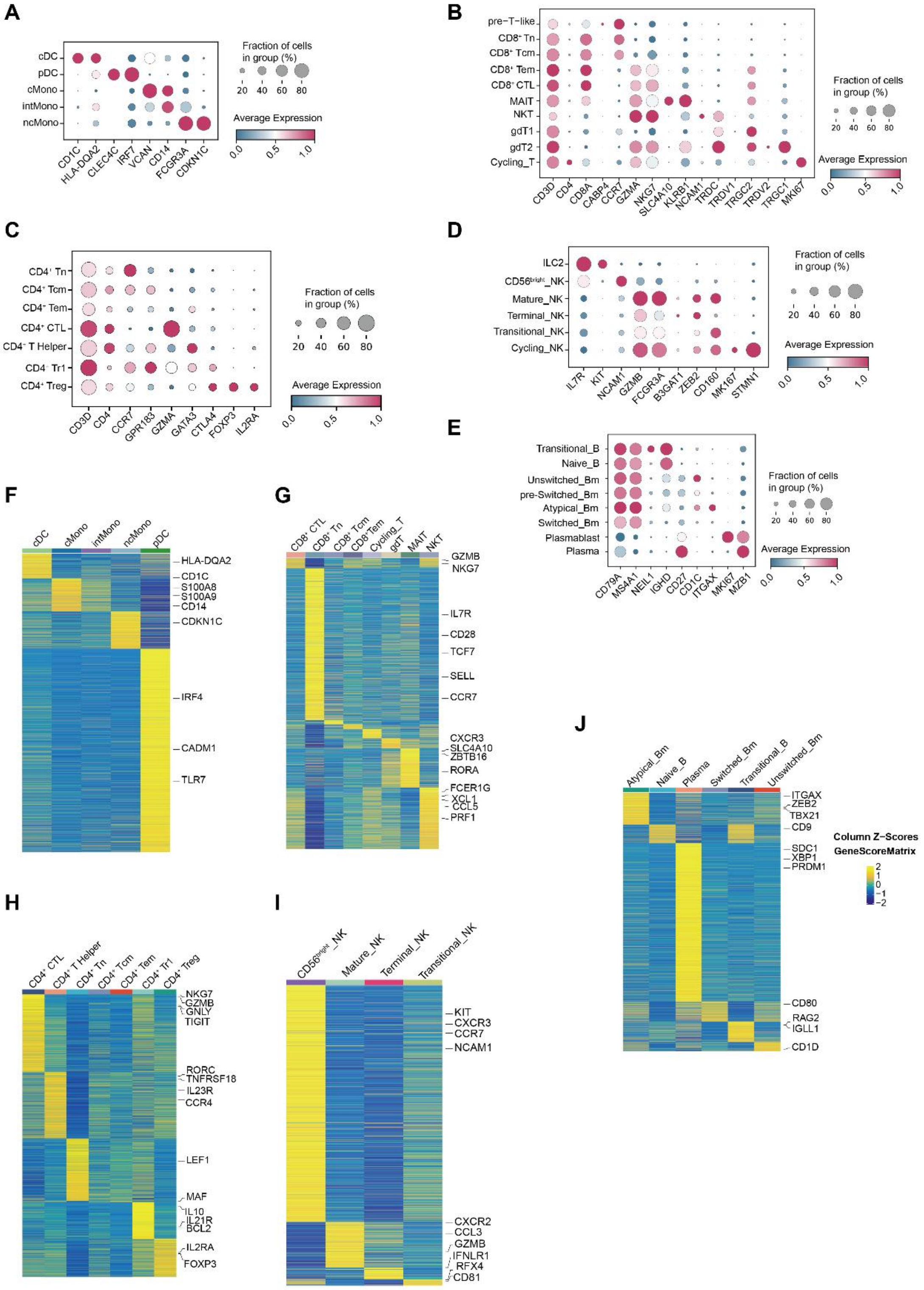
A
cDC
pDC
cMono
intMono
ncMono
CD1C
HLA-DQA2
CLEC4C
IRF7
VCAN
CD14
FCGR3A
CDKN1C
Fraction of cells in group (%)
20 40 60 80
Average Expression
0.0 0.5 1.0
B
pre−T−like
CD8⁺ Tn
CD8⁺ Tcm
CD8⁺ Tem
CD8⁺ CTL
MAIT
NKT
gdT1
gdT2
Cycling_T
CD3D
CD4
CD8A
CABP4
CCR7
GZMA
NKG7
SLC4A10
KLRB1
NCAM1
TRDC
TRDV1
TRGC2
TRDV2
TRGC1
MKI67
Fraction of cells in group (%)
20 40 60 80
Average Expression
0.0 0.5 1.0
C
CD4⁺ Tn
CD4⁺ Tcm
CD4⁺ Tem
CD4⁺ CTL
CD4⁺ T Helper
CD4⁺ Tr1
CD4⁺ Treg
CD3D
CD4
CCR7
GPR183
GZMA
GATA3
CTLA4
FOXP3
IL2RA
Fraction of cells in group (%)
20 40 60 80
Average Expression
0.0 0.5 1.0
D
ILC2
CD56bright_NK
Mature_NK
Terminal_NK
Transitional_NK
Cycling_NK
IL7R
KIT
NCAM1
GZMB
FCGR3A
B3GAT1
ZEB2
CD160
MKI67
STMN1
Fraction of cells in group (%)
20 40 60 80
Average Expression
0.0 0.5 1.0
E
Transitional_B
Naive_B
Unswitched_Bm
pre-Switched_Bm
Atypical_Bm
Switched_Bm
Plasmablast
Plasma
CD79A
MS4A1
NEIL1
IGHD
CD27
CD1C
ITGAX
MKI67
MZB1
Fraction of cells in group (%)
20 40 60 80
Average Expression
0.0 0.5 1.0
F
cDC
cMono
intMono
ncMono
pDC
HLA-DQA2
CD1C
S100A8
S100A9
CD14
CDKN1C
IRF4
CADM1
TLR7
G
CD8⁺ CTL
CD8⁺ Tn
CD8⁺ Tcm
CD8⁺Tem
Cycling_T
gdT
MAIT
NKT
GZMB
NKG7
IL7R
CD28
TCF7
SELL
CCR7
CXCR3
SLC4A10
ZBTB16
RORA
FCER1G
XCL1
CCL5
PRF1
H
CD4⁺ CTL
CD4⁺ T Helper
CD4⁺ Tn
CD4⁺ Tcm
CD4⁺ Tem
CD4⁺ Tr1
CD4⁺ Treg
NKG7
GZMB
GNLY
TIGIT
RORC
TNFRSF18
IL23R
CCR4
LEF1
MAF
IL10
IL21R
BCL2
IL2RA
FOXP3
I
CD56bright_NK
Mature_NK
Terminal_NK
Transitional_NK
KIT
CXCR3
CCR7
NCAM1
CXCR2
CCL3
GZMB
IFNLR1
RFX4
CD81
J
Atypical_Bm
Naive_B
Plasma
Switched_Bm
Transitional_B
Unswitched_Bm
ITGAX
ZEB2
TBX21
CD9
SDC1
XBP1
PRDM1
CD80
RAG2
IGLL1
CD1D
Column Z-Scores
GeneScoreMatrix
2
1
0
−1
−2

### Figure S5

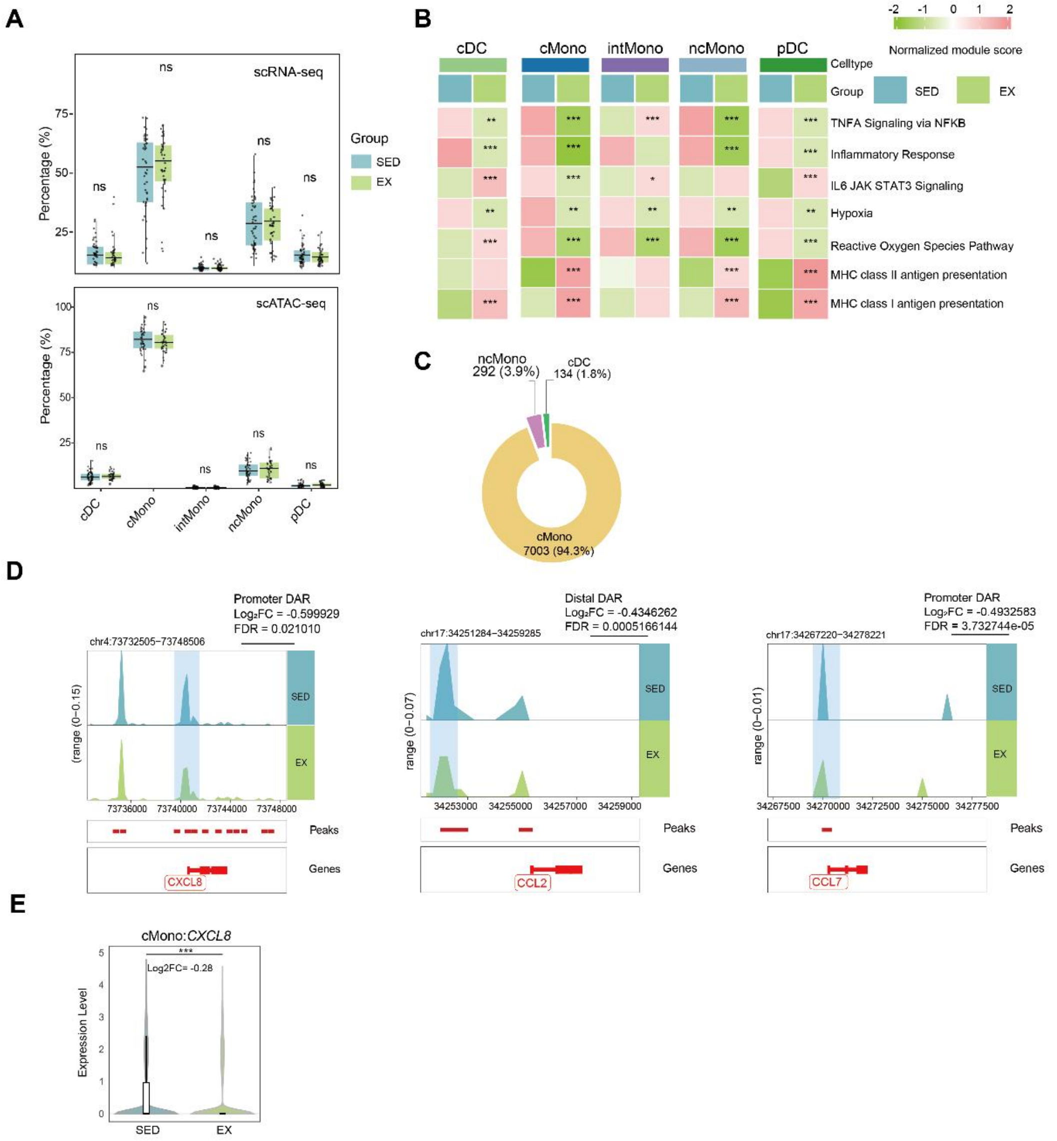

A
B
C
D
E
scRNA-seq
scATAC-seq
Percentage (%)
Group
SED
EX
cDC
cMono
intMono
ncMono
pDC
Normalized module score
Celltype
TNFA Signaling via NFKB
Inflammatory Response
IL6 JAK STAT3 Signaling
Hypoxia
Reactive Oxygen Species Pathway
MHC class II antigen presentation
MHC class I antigen presentation
ncMono
292 (3.9%)
cDC
134 (1.8%)
cMono
7003 (94.3%)
Promoter DAR
Log2FC = -0.599929
FDR = 0.021010
chr4:73732505-73748506
Distal DAR
Log2FC = -0.4346262
FDR = 0.0005166144
chr17:34251284-34259285
Promoter DAR
Log2FC = -0.4932583
FDR = 3.732744e-05
chr17:34267220-34278221
Peaks
Genes
CXCL8
CCL2
CCL7
cMono:CXCL8
Log2FC= -0.28
Expression Level

### Figure S6

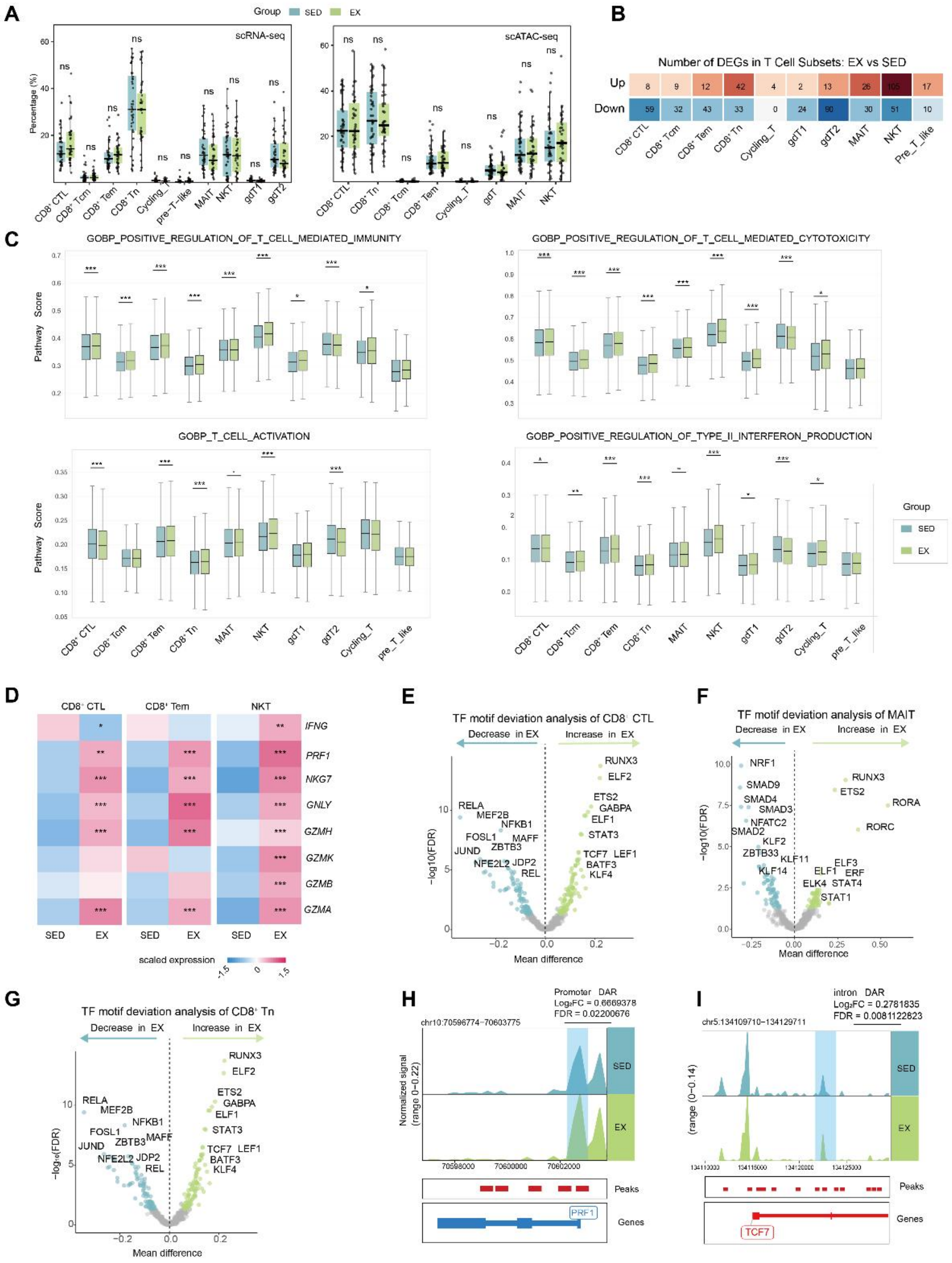

A
B
Number of DEGs in T Cell Subsets: EX vs SED
C
GOBP_POSITIVE_REGULATION_OF_T_CELL_MEDIATED_IMMUNITY
GOBP_POSITIVE_REGULATION_OF_T_CELL_MEDIATED_CYTOTOXICITY
GOBP_T_CELL_ACTIVATION
GOBP_POSITIVE_REGULATION_OF_TYPE_II_INTERFERON_PRODUCTION
D
E
TF motif deviation analysis of CD8+ CTL
F
TF motif deviation analysis of MAIT
G
TF motif deviation analysis of CD8+ Tn
H
I

### Figure S7

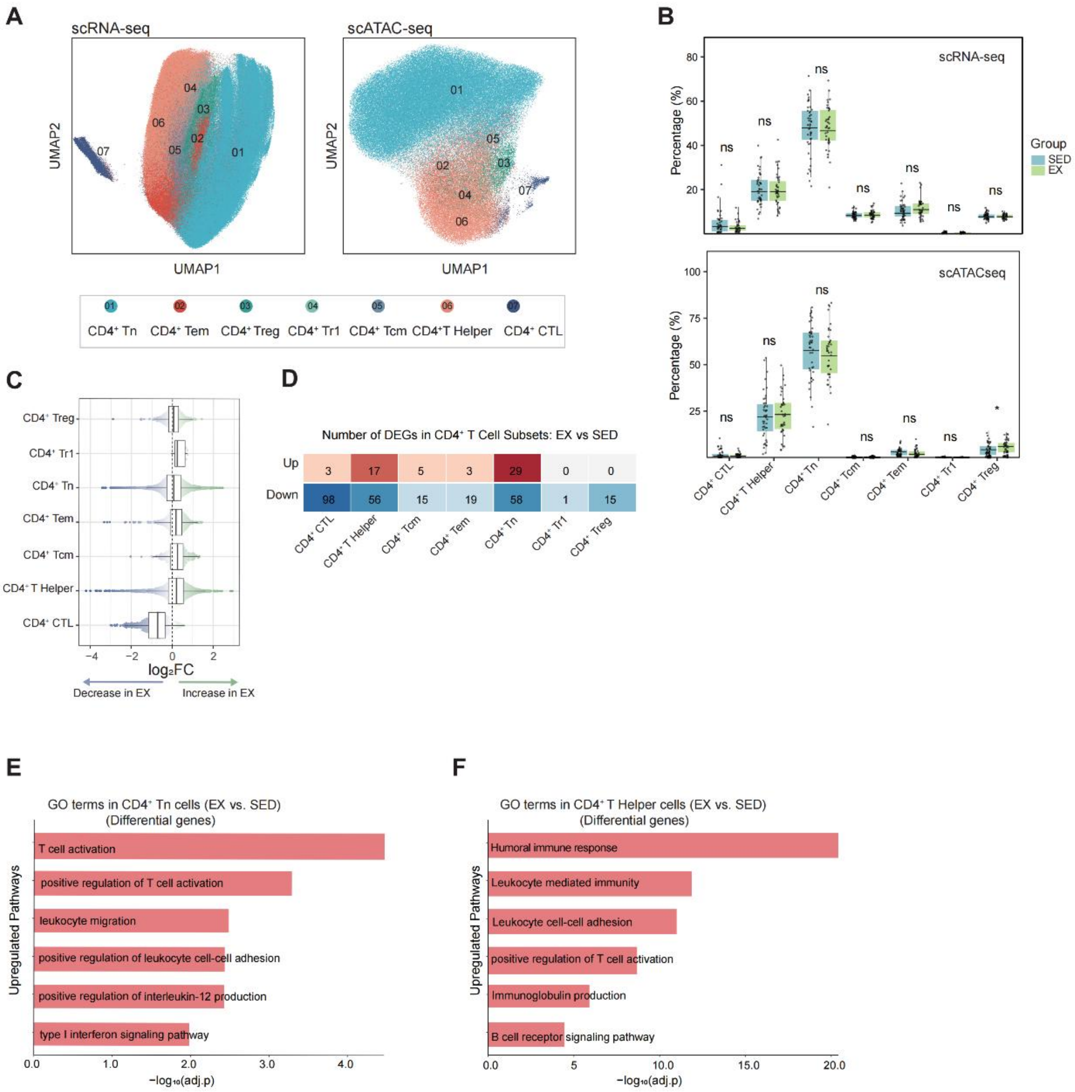
A
scRNA-seq
scATAC-seq
UMAP1
UMAP2
01
02
03
04
05
06
07
CD4⁺ Tn
CD4⁺ Tem
CD4⁺ Treg
CD4⁺ Tr1
CD4⁺ Tcm
CD4⁺T Helper
CD4⁺ CTL
B
scRNA-seq
scATACseq
Percentage (%)
ns
*
Group
SED
EX
CD4⁺ CTL
CD4⁺T Helper
CD4⁺ Tn
CD4⁺ Tcm
CD4⁺ Tem
CD4⁺ Tr1
CD4⁺ Treg
C
CD4⁺ Treg
CD4⁺ Tr1
CD4⁺ Tn
CD4⁺ Tem
CD4⁺ Tcm
CD4⁺ T Helper
CD4⁺ CTL
log₂FC
Decrease in EX
Increase in EX
D
Number of DEGs in CD4⁺ T Cell Subsets: EX vs SED
Up
3
17
5
3
29
0
0
Down
98
56
15
19
58
1
15
CD4⁺ CTL
CD4⁺ T Helper
CD4⁺ Tcm
CD4⁺ Tem
CD4⁺ Tn
CD4⁺ Tr1
CD4⁺ Treg
E
GO terms in CD4⁺ Tn cells (EX vs. SED)
(Differential genes)
Upregulated Pathways
T cell activation
positive regulation of T cell activation
leukocyte migration
positive regulation of leukocyte cell-cell adhesion
positive regulation of interleukin-12 production
type I interferon signaling pathway
−log₁₀(adj.p)
F
GO terms in CD4⁺ T Helper cells (EX vs. SED)
(Differential genes)
Upregulated Pathways
Humoral immune response
Leukocyte mediated immunity
Leukocyte cell-cell adhesion
positive regulation of T cell activation
Immunoglobulin production
B cell receptor signaling pathway
−log₁₀(adj.p)

### Figure S8

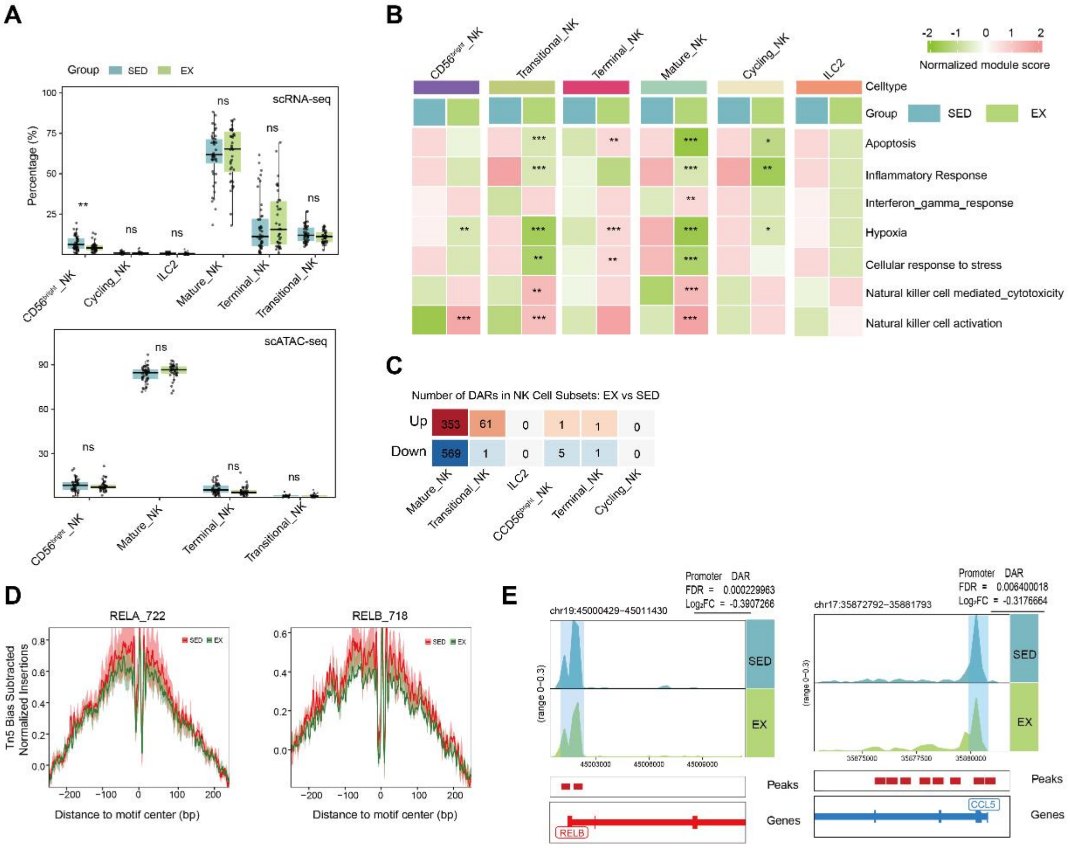

A
Group
SED
EX
scRNA-seq
Percentage (%)
scATAC-seq
CD56bright_NK
Cycling_NK
ILC2
Mature_NK
Terminal_NK
Transitional_NK
B
Normalized module score
Celltype
Group
Apoptosis
Inflammatory Response
Interferon_gamma_response
Hypoxia
Cellular response to stress
Natural killer cell mediated_cytotoxicity
Natural killer cell activation
C
Number of DARs in NK Cell Subsets: EX vs SED
Up
Down
353
61
0
1
1
0
569
1
0
5
1
0
D
RELA_722
RELB_718
Tn5 Bias Subtracted Normalized Insertions
Distance to motif center (bp)
E
chr19:45000429-45011430
Promoter DAR
FDR = 0.000229963
Log2FC = -0.3807266
chr17:35872792-35881793
FDR = 0.006400018
Log2FC = -0.3176664
Peaks
Genes
RELB
CCL5

#### Figure S9

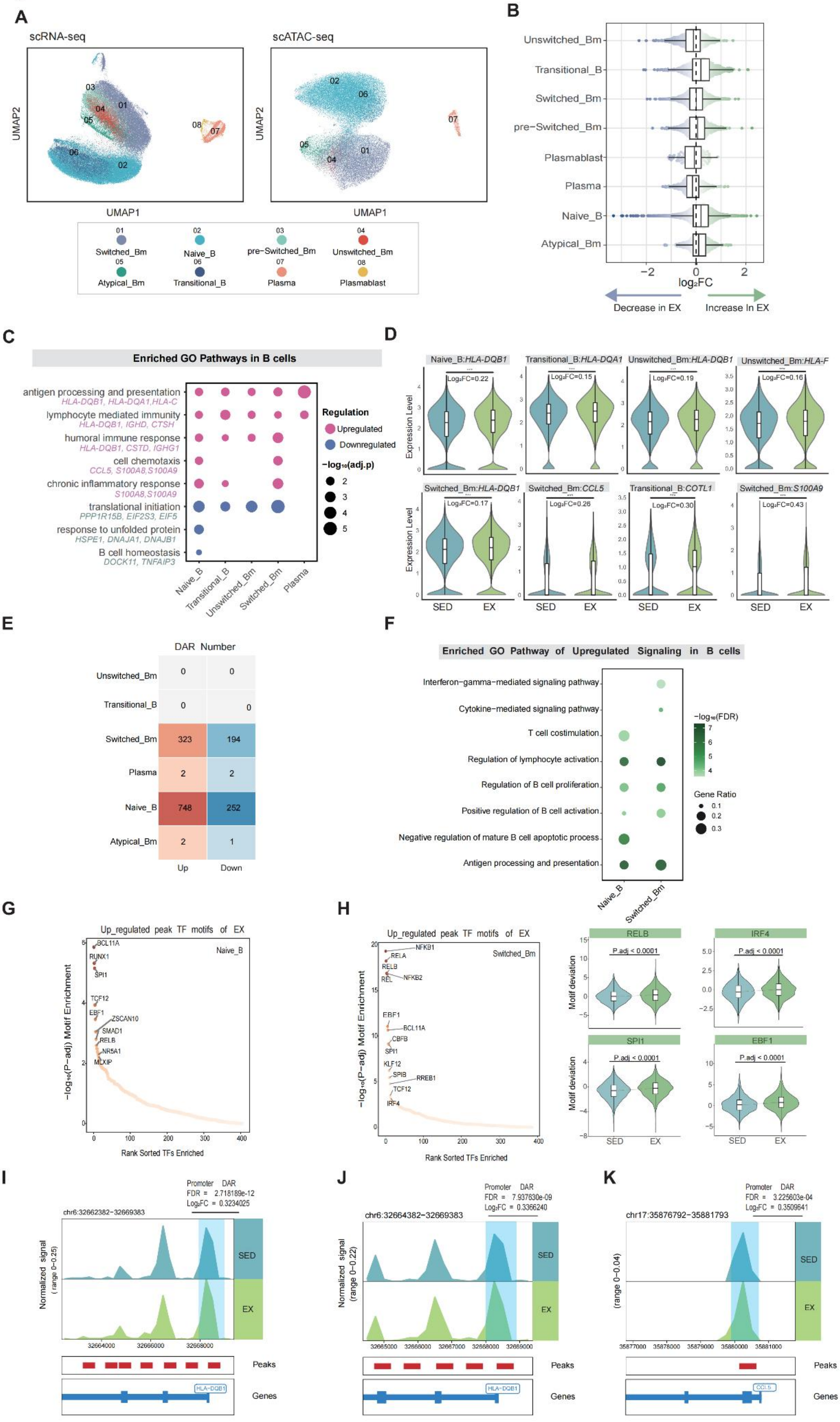
A
scRNA-seq
scATAC-seq
UMAP1
UMAP2
01 Switched_Bm
02 Naive_B
03 pre−Switched_Bm
04 Unswitched_Bm
05 Atypical_Bm
06 Transitional_B
07 Plasma
08 Plasmablast
B
Unswitched_Bm
Transitional_B
Switched_Bm
pre−Switched_Bm
Plasmablast
Plasma
Naive_B
Atypical_Bm
log₂FC
Decrease in EX
Increase In EX
C
Enriched GO Pathways in B cells
antigen processing and presentation
HLA-DQB1, HLA-DQA1,HLA-C
lymphocyte mediated immunity
HLA-DQB1, IGHD, CTSH
humoral immune response
HLA-DQB1, CSTD, IGHG1
cell chemotaxis
CCL5, S100A8,S100A9
chronic inflammatory response
S100A8,S100A9
translational initiation
PPP1R15B, EIF2S3, EIF5
response to unfolded protein
HSPE1, DNAJA1, DNAJB1
B cell homeostasis
DOCK11, TNFAIP3
Regulation
Upregulated
Downregulated
−log₁₀(adj.p)
Naive_B
Transitional_B
Unswitched_Bm
Switched_Bm
Plasma
D
Naive_B:HLA-DQB1
Transitional_B:HLA-DQA1
Unswitched_Bm:HLA-DQB1
Unswitched_Bm:HLA-F
Switched_Bm:HLA-DQB1
Switched_Bm:CCL5
Transitional_B:COTL1
Switched_Bm:S100A9
Expression Level
Log₂FC=0.22
Log₂FC=0.15
Log₂FC=0.19
Log₂FC=0.16
Log₂FC=0.17
Log₂FC=0.26
Log₂FC=0.30
Log₂FC=0.43
SED
EX
E
DAR Number
Unswitched_Bm 0 0
Transitional_B 0 0
Switched_Bm 323 194
Plasma 2 2
Naive_B 748 252
Atypical_Bm 2 1
Up
Down
F
Enriched GO Pathway of Upregulated Signaling in B cells
Interferon−gamma−mediated signaling pathway
Cytokine−mediated signaling pathway
T cell costimulation
Regulation of lymphocyte activation
Regulation of B cell proliferation
Positive regulation of B cell activation
Negative regulation of mature B cell apoptotic process
Antigen processing and presentation
−log₁₀(FDR)
Gene Ratio
Naive_B
Switched_Bm
G
Up_regulated peak TF motifs of EX
Naive_B
BCL11A
RUNX1
SPI1
TCF12
EBF1
ZSCAN10
SMAD1
RELB
NR5A1
MLXIP
−log₁₀(P−adj) Motif Enrichment
Rank Sorted TFs Enriched
H
Up_regulated peak TF motifs of EX
Switched_Bm
NFKB1
RELA
RELB
REL
NFKB2
EBF1
BCL11A
CBFB
SPI1
KLF12
SPIB
RREB1
TCF12
IRF4
−log₁₀(P−adj) Motif Enrichment
Rank Sorted TFs Enriched
RELB
IRF4
SPI1
EBF1
P.adj < 0.0001
Motif deviation
SED
EX
I
Promoter DAR
FDR = 2.718189e-12
Log₂FC = 0.3234025
chr6:32662382−32669383
Normalized signal (range 0−0.25)
SED
EX
Peaks
Genes
HLA-DQB1
J
Promoter DAR
FDR = 7.937630e-09
Log₂FC = 0.3366240
chr6:32664382−32669383
Normalized signal (range 0−0.22)
SED
EX
Peaks
Genes
HLA-DQB1
K
Promoter DAR
FDR = 3.225603e-04
Log₂FC = 0.3509641
chr17:35876792−35881793
(range 0−0.04)
SED
EX
Peaks
Genes
CCL5

### Figure S10

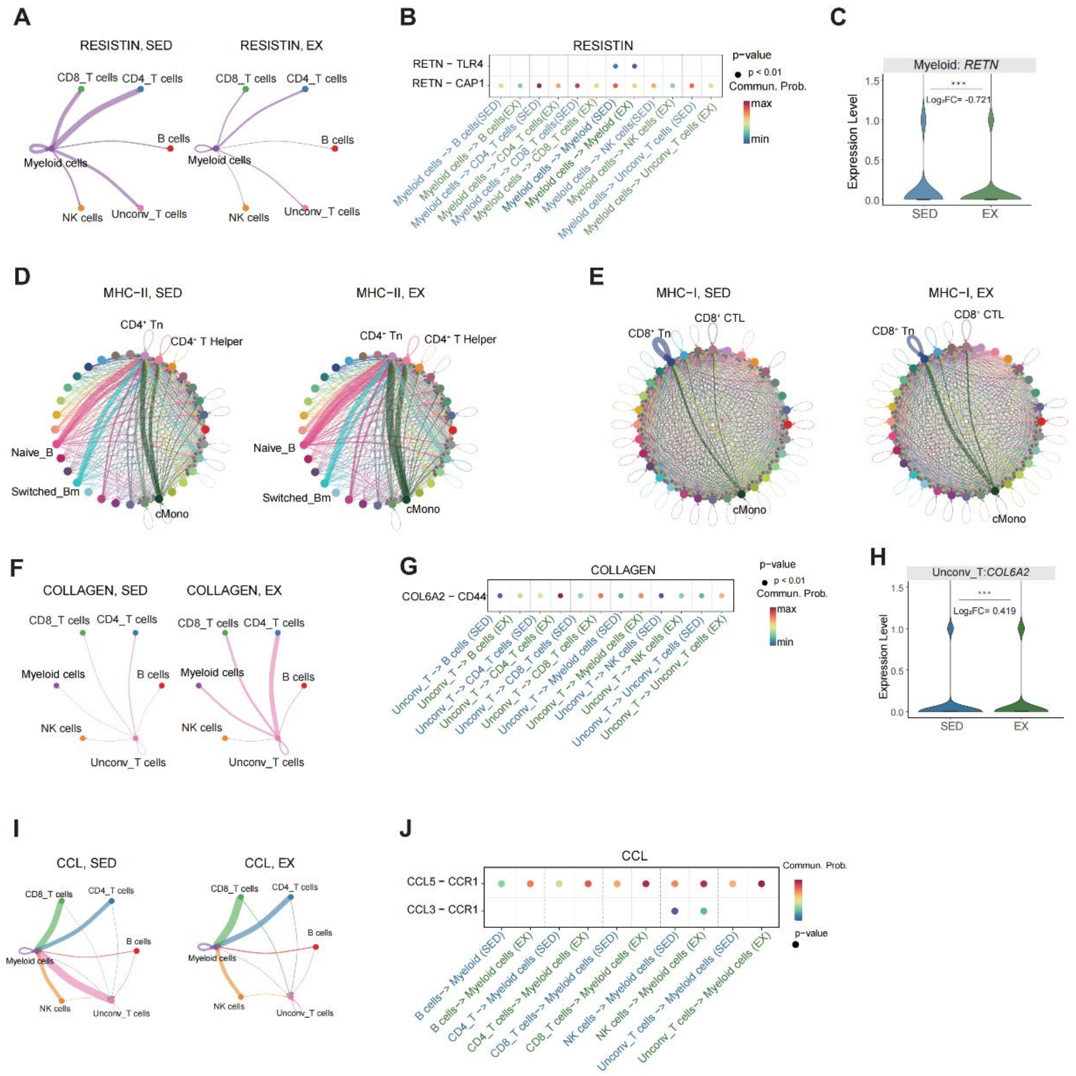
A
RESISTIN, SED
RESISTIN, EX
CD8_T cells
CD4_T cells
B cells
Myeloid cells
NK cells
Unconv_T cells
B
RESISTIN
RETN - TLR4
RETN - CAP1
p-value
p < 0.01
Commun. Prob.
max
min
C
Myeloid: RETN
Expression Level
Log2FC= -0.721
SED
EX
D
MHC-II, SED
MHC-II, EX
CD4+ Tn
CD4+ T Helper
Naive_B
Switched_Bm
cMono
E
MHC-I, SED
MHC-I, EX
CD8+ Tn
CD8+ CTL
cMono
F
COLLAGEN, SED
COLLAGEN, EX
G
COLLAGEN
COL6A2 - CD44
H
Unconv_T:COL6A2
Log2FC= 0.419
I
CCL, SED
CCL, EX
J
CCL
CCL5 - CCR1
CCL3 - CCR1
